# Phenomic Prediction I: Plot-Level Prediction of Lodging Severity in Sorghum Breeding Trials Using UAV-Based Photogrammetric Height Data

**DOI:** 10.64898/2026.03.26.713817

**Authors:** Srinivasa Reddy Mothukuri, Sean Reynolds Massey-Reed, Andries Potgieter, Kenneth Laws, Colleen Hunt, Esinam Nancy Amuzu-Aweh, Mark Cooper, Emma Mace, David Jordan

## Abstract

Lodging in sorghum presents a significant challenge for plant breeders due to the trade-off between lodging resistance and grain yield. Manually measuring lodging across thousands of plots is time-consuming, expensive, and error-prone, making selection for lodging resistance challenging in breeding programs. Unmanned aerial vehicle (UAV)-derived metrics provide a potential high-throughput alternative; however, it remains unclear whether photogrammetric heights derived from UAV imagery can estimate plot-level lodging severity in large sorghum breeding trials.

This study developed a framework for predicting plot-level lodging from UAV imagery across 2,675 sorghum breeding plots. Multi-temporal canopy height data were collected at two critical time points: maximum crop height and at manual lodging assessment. Height percentiles were extracted from UAV-derived point clouds generated using photogrammetric algorithms. These data were used to develop parametric, non-parametric, and ensemble prediction models, which were evaluated using three statistical metrics.

The ensemble model, averaging predictions from all models, achieved the highest accuracy with Pearson correlations of r = 0.80-0.84 and lowest root mean square error (RMSE=16-18%), explaining 64-70% of variation in manual lodging counts. Model diagnostics and iterative refinement, including inspection of UAV imagery and dataset curation, had minimal impact on model performance, demonstrating the robustness of the approach. Model performance was consistent across sites, with minimal effects of stratified sampling on accuracy, confirming the ensemble approach as optimal for plot-level lodging assessment.

This study demonstrates that integrated multi-temporal UAV imagery offers a practical alternative to labor-intensive manual evaluation methods by enabling high-throughput lodging assessment suitable for implementation in sorghum breeding programs.

**PLAIN LANGUAGE SUMMARY:** Lodging happens when sorghum plants fall over before harvest, which can reduce grain yield and make harvesting difficult. Breeders need to measure lodging in large field trials, but manually scoring thousands of plots takes a lot of time, is expensive, and can give different results depending on who does the scoring. This study tested whether height information from drone images could be used to estimate lodging in sorghum breeding plots. The drone-based measurements matched well with manual lodging counts, and the results were consistent across different sites. This shows that drone imagery can be a faster and more affordable way to measure lodging and help breeders select for lodging resistance in sorghum breeding programs.

## 1. INTRODUCTION

Lodging is the permanent displacement of crop stems from their vertical position. It can be caused either by the breaking of the stem (stem lodging) or by root rotation in the soil (root lodging) (Pinthus 1974). Stem lodging is the predominant type of lodging in sorghum and is widely observed in countries where hybrid sorghum is grown, such as Australia (Henzell et al. 1984), the United States of America (USA) (Rosenow 1977), and Argentina (Frezzi & Teyssandier 1980). Sorghum is typically grown as a rain-fed crop in different water-limited environments, and as a result sorghum crops frequently encounter water stress at various stages of development, with drought during the grain-filling stage being common and having significant impacts on grain yield (Chapman et al. 2000). In addition to directly reducing yield, water stress during grain filling reduces photosynthesis, causing the remobilisation of carbon from the stem to the panicle, which weakens the stem and can result in stem lodging (Borrell & Douglas 1996).

Sorghum plants with high harvest index, typically observed in hybrids, have a larger demand for carbohydrates compared to supply and tend to be more prone to lodging. This balance between supply and demand sets up a trade-off between lodging resistance and grain yield potential (Alam et al. 2014), with hybrids with high yield potential tending to be more susceptible to lodging (Wang et al. 2020). The negative relationship between these two key traits creates a major challenge for plant breeders who are attempting to produce high-yielding hybrids that do not lodge.

In sorghum breeding programs, lodging is typically quantified through visual scores. In visual scoring, lodging is assessed by visually inspecting the plot and assigning a score between 1 and 9 based on the severity of lodging. A score of 1 indicates 0 to 11% lodging, while a score of 9 represents 89 to 100% lodging in a plot. Visual scoring is time-consuming and prone to human error and biases (Chapman et al. 2014; Hu et al. 2018; Potgieter et al. 2017), which can negatively impact breeding efficiency and the accurate identification of optimal genotypes (Chapman et al. 2014). Physical counting of lodged stems is a more accurate method of assessing lodging but is highly labour intensive which precludes its use in typical plant breeding programs, which need to evaluate large numbers of entries at multiple sites. As a result, there is a significant need to develop high throughput approaches for estimating lodging, such as using unmanned aerial vehicle (UAV) derived photogrammetric methods.

Sensors attached to UAVs, capturing images, offer a more cost-effective and flexible method for data collection. UAVs can be fitted with a wide variety of sensors, allowing them to gather high-resolution spatial and temporal data quickly, cheaply and non-destructively (Chapman et al. 2014; Potgieter et al. 2017; Zhao et al. 2021). This technology enables rapid phenotyping of entire fields by following pre-programmed flight paths with specific altitudes and speeds, thereby greatly improving the efficiency and scalability of large-scale field research (Chapman et al. 2014; Hu et al. 2018; Potgieter et al. 2017). Recently, there have been numerous examples demonstrating how UAVs equipped with various sensors such as RGB, multispectral, and hyperspectral can efficiently gather diverse remote sensing information in a single flight (Furbank et al. 2019; Furbank & Tester 2011; Zhi et al. 2022).

In recent years, UAVs have become increasingly popular for collecting canopy height data (Hu et al. 2018) due to its importance as a growth indicator closely associated with canopy coverage (Lee & Lee 2013), lodging resistance, biomass, and yield performance (Gao et al. 2020; D. Wang et al. 2022; Zhi et al. 2022). Various algorithms have been utilized to predict agronomic traits from sensor-derived metrics, enabling significant advancements in plant breeding and agriculture. Applying machine learning techniques for this purpose remains a current and significant area of research in sensor-derived metrics, offering a powerful and efficient method for estimating various agronomic traits (Ashapure et al. 2020; Dang et al. 2021; Smith et al. 2024). The statistical and machine learning algorithms have the ability to analyse diverse datasets through both parametric and nonparametric methods, including UAV-based photogrammetry method.

Parametric models such as multiple linear regression (MLR), ridge regression (RR), and partial least squares (PLS) are commonly used in UAV-based (image/spectral) estimation of crop canopy traits related to biomass, leaf area index (LAI) or Photosynthetic Capacity (Vcmax) (Sun et al. 2020; Usai et al. 2009; Zhi et al. 2022) and assume a uniform response across all conditions. In contrast, non-parametric models, which do not assume a linear relationship between variables, include algorithms like support vector machines radial (SVM Radial), K-nearest neighbours (KNN), and random forest (RF) for capturing complex, non-linear data relationships (Breiman 2001). Non-parametric models often require significant computation, including parameter tuning, which impacts robustness. Processing large UAV datasets also demands lengthy training times and high processing power. These parametric and non-parametric machine-learning algorithms enhance and accelerate the identification of relationships among plant phenotypic traits, such as plant height, biomass, and yield (Ashapure et al. 2020; Dang et al. 2021). An alternative strategy involves exploring combinations of diverse individual predictive models using ensemble learning to improve performance and robustness.

In plant breeding, no single predictive model has provided the best prediction accuracy across all traits, trials, and populations. This principle is consistent with the ‘No Free Lunch Theorem’ proposed by (Wolpert & Macready 1997), which clarifies that the relative performance of any algorithm depends on the characteristics of the prediction problem. One method of exploiting this variability in model performance is to select the superior model for each scenario. A robust starting point is to exploit the complementary strengths of different models by combining them in an ensemble-based approach (Page 2018; Tomura et al. 2025). Ensemble learning, a specialized area within machine learning, has emerged as an important approach for estimating agronomical traits in recent years. By leveraging the combined strengths of multiple models, ensemble methods generally outperform individual learners in predictive accuracy (Tomura et al. 2025; Zhou 2012). Researchers have highlighted the substantial promise of ensemble learning for applications in plant phenotyping. For instance, wheat yield predictions across varying treatments were effectively achieved by integrating UAV-derived multispectral metrics with ensemble learnings (Fei et al. 2021).

Previous studies have applied different UAV-based approaches to assess crop lodging. Chu et al. (2017) used UAV-based Structure-from-Motion photogrammetry to generate canopy height models in maize and estimated lodging severity by classifying individual row segments as lodged or non-lodged using the 90^th^ and 99^th^ percentile canopy heights and predefined height thresholds. In addition, L. Zhou et al. (2020) employed UAV-LiDAR data to quantify plant-height changes in artificially lodged maize and evaluated the recovery of different lodging types over time. Shu et al. (2023) investigated changes in canopy colour and image texture variables derived from UAV images collected before and after lodging and used regression and machine-learning models to classify maize into four lodging severity categories. In wheat, Zhang et al. (2020) combined RGB and multispectral UAV images with transfer learning and a DeepLabv3+ deep-learning model to automatically identify wheat lodging areas across multiple crop growth stages. Sarkar et al. (2023) examined differences in the spatial pattern and uniformity of soybean canopies derived from UAV RGB images, together with machine-learning models, to classify plots into four lodging severity categories. Li et al. (2024) derived vegetation indices derived from UAV RGB and multispectral imagery at different crop growth stages, together with machine-learning models, to predict stem and root lodging susceptibility in rapeseed. A recent study by Zhou et al. (2026) extracted deep spatial features from UAV multispectral imagery using YOLOv8, YOLO12, and Segment Anything models and combined these features with machine-learning classifiers to distinguish lodged and non-lodged areas in winter wheat.

However, it remains unclear whether photogrammetry-derived percentile heights from UAV imagery can be used to predict plot-level lodging percentages in large sorghum breeding trials. Specifically, we aimed to: (i) identify UAV-derived predictors that effectively capture lodging severity; (ii) develop and evaluate predictive models using a cross-validation (resampling) approach; and (iii) assess the factors influencing prediction accuracy.

## 2. MATERIALS AND METHODS

### 2.1 Trials

This study utilized existing data from trials conducted at two breeding trial locations (Jandowae and Pirrinuan) in Queensland during the 2018-2019 growing season. The Pirrinuan site is located at 27.096°S latitude and 151.222°E longitude, with an elevation of 356 meters above sea level. The Jandowae site is located at 26.891°S latitude and 151.128°E longitude, with an elevation of 344 meters above sea level. Each site contained two sorghum hybrid trial series designated as advanced yield trial for males (AYTM) and advanced yield trial for females (AYTF). The AYTM trial contained a set of advanced candidate male parents crossed with two female testers and the AYTF trial contained a set of advanced candidate female parents crossed with two male testers. In addition, the trials contained a range of commercial check hybrids.

These trials were part of the sorghum pre-breeding program conducted by the University of Queensland (UQ), the Department of Primary Industries (DPI), and the Grains Research and Development Corporation (GRDC). The trials were managed with standard agronomic practices for the region. The trial series were designed with partial replication (Cullis et al. 2006), where some genotypes were replicated, and others had a single replicate, incorporating checks. Genotypes were arranged using a spatial row-column design, with replicated hybrids divided into two equal blocks. However, our research focused on plot-based analysis without considering genotype or spatial effects. The planting dates and numbers of entries and plots are described in Table 1. Since the primary purpose of this research was to estimate plot lodging rather than genotype performance the trial designs and genotypes were not used in subsequent analyses.

**Table 1:** Summary of the manual count and UAV sensing data by site, and trial series for genotypes and plots. AYTF: advanced yield trial female; AYTM: advanced yield trial male. Flight-0 was conducted at crop emergence, Flight-1 around the flowering stage, and Flight-2 on the day lodging was measured.

| Site | Trial | Plots | Sowing | Flight-0 | Flight-1 | Flight-2 |
| --- | --- | --- | --- | --- | --- | --- |
| Jandowae | AYTF | 750 | 1/11/2018 | 13/11/2018 | 16/01/2019 | 28/02/2019 |
|  | AYTM | 600 | 1/11/2018 | 13/11/2018 | 16/01/2019 | 28/02/2019 |
| Pirrinuan | AYTF | 750 | 1/11/2018 | 13/11/2018 | 16/01/2019 | 01/03/2019 |
|  | AYTM | 600 | 1/11/2018 | 13/11/2018 | 16/01/2019 | 01/03/2019 |

### 2.2 Manual lodging counts and UAV data collection

Manual lodging counts were collected for all plots at Jandowae and Pirrinuan (Table 1). Each site included two trials. In three of the four trial series (trial by site), the total number of plants and the number of lodged plants were counted individually within each plot. Manual lodging percentage was then calculated using Equation (1).

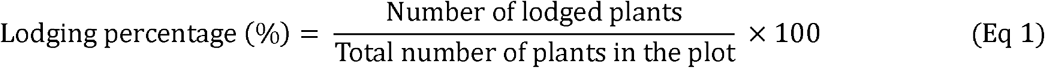

Values ranged from 0%, indicating that no plants in the plot were lodged, to 100%, indicating that all plants were lodged. In the Pirrinuan AYTM trial, a modified counting method was used because the time available for lodging assessment was limited due to approaching adverse weather. Lodging was therefore assessed using approximate counts of lodged plants relative to the total number of plants within each plot, and the resulting lodging percentages were recorded in increments of 5 percentage points (i.e., 0, 5, 10, 15, …, 95, and 100%).

To maintain a consistent recording scale across all four-trial series, lodging percentages from the other trials were subsequently rounded to the nearest multiple of 5 before analysis.

UAV flights were conducted across the entire field and all trial series on three dates for each trial (Table 1).

### 2.3 UAV imagery pre-processing

The approach proposed here uses a minimum of three UAV flights to make this approach practically applicable in a plant breeding program. Flight-0 was conducted at crop emergence to quantify establishment and derive ground (bare-soil) reference heights. Flight-1 was conducted at the flowering stage (approximately 76 days after sowing), when the crop approached maximum canopy height. Flight-2 was conducted on the day lodging was assessed (approximately one week before harvest or ∼120 days after sowing).

The UAV imagery pre-processing workflow begins with precise ground control point (GCP) placement and measurement using global navigation satellite system (GNSS) technology capable of sub-1 cm positioning (Fig S1). The UAV was equipped with a multispectral sensor and flown at an altitude of approximately 30 meters for flight 0 and flight 1 and 35 meters for flight 2 over the trial plots (Fig S1). The UAV captured 16-bit TIF images and calibration images, which were systematically downloaded to a network attached storage (NAS) system and labelled. The image processing pipeline utilized Pix4D (photogrammetry) software (Fig S1), initiating with an initial calibration step to align image positions and generate preliminary JPG previews from the original TIF files.

GCP locations were imported and marked in each image, ensuring accurate georeferencing of the project. This process allowed for accurate height calculations and maintained consistent spatial alignment across multiple flight sessions. After re-running the initial processing step with the georeferencing data, the software generated point clouds through a densification process, first processing them in separate blocks before merging them into a single LAS format file. The full-resolution LAS file was subsequently downscaled to a 2.5D 2 cm resolution (Fig S1). At the same time, a shapefile was generated to define individual plot boundaries, assigning each a unique plot identifier. The processed data, including reflectance maps, shapefiles, and LAS files, was systematically organized into a structured directory. A custom Python script was used to run the extraction program (Xtractori) (Das et al. 2022), which generated a CSV file containing reflected pixel values and height data at the plot level (Fig S1). A final python script then reformatted the CSV, making it more flexible for analysis.

#### 2.3.1 Percentile height index (PHI)

The PHI for each plot was determined by analysing three-dimensional point cloud data from UAV imagery. Height measurements were obtained directly from the LAS 2.5 DSM grid files, with the focus on calculating height percentiles for each plot, excluding any vegetation indices. The analysis aimed to provide a detailed representation of the height distribution by calculating percentile values from the 1^st^ to the 99^th^ percentiles.

The methodology used to determine canopy height from the point cloud data involved accounting for variations in elevation across the field. Height values from the 1^st^ to 99^th^ percentiles were calculated (Fig 1) after adjusting for elevation above sea level on a plot-by-plot basis, as field slopes typically vary across fields but remain relatively minimal within individual plots. For each plot, canopy height was calculated by taking into account the median height of that plot at emergence. The median emergence height was used as the reference height because the soil surface was mostly visible at this stage and provided a stable plot-level baseline. In high-stubble fields, the 2^nd^ percentile height at emergence was used instead of the median because standing stubble could increase the median height and reduce its suitability as a ground reference. The 2^nd^ percentile was selected because it better represented the lower surface of the plot while avoiding extreme low values that may occur from point-cloud noise.

**Fig 1:**
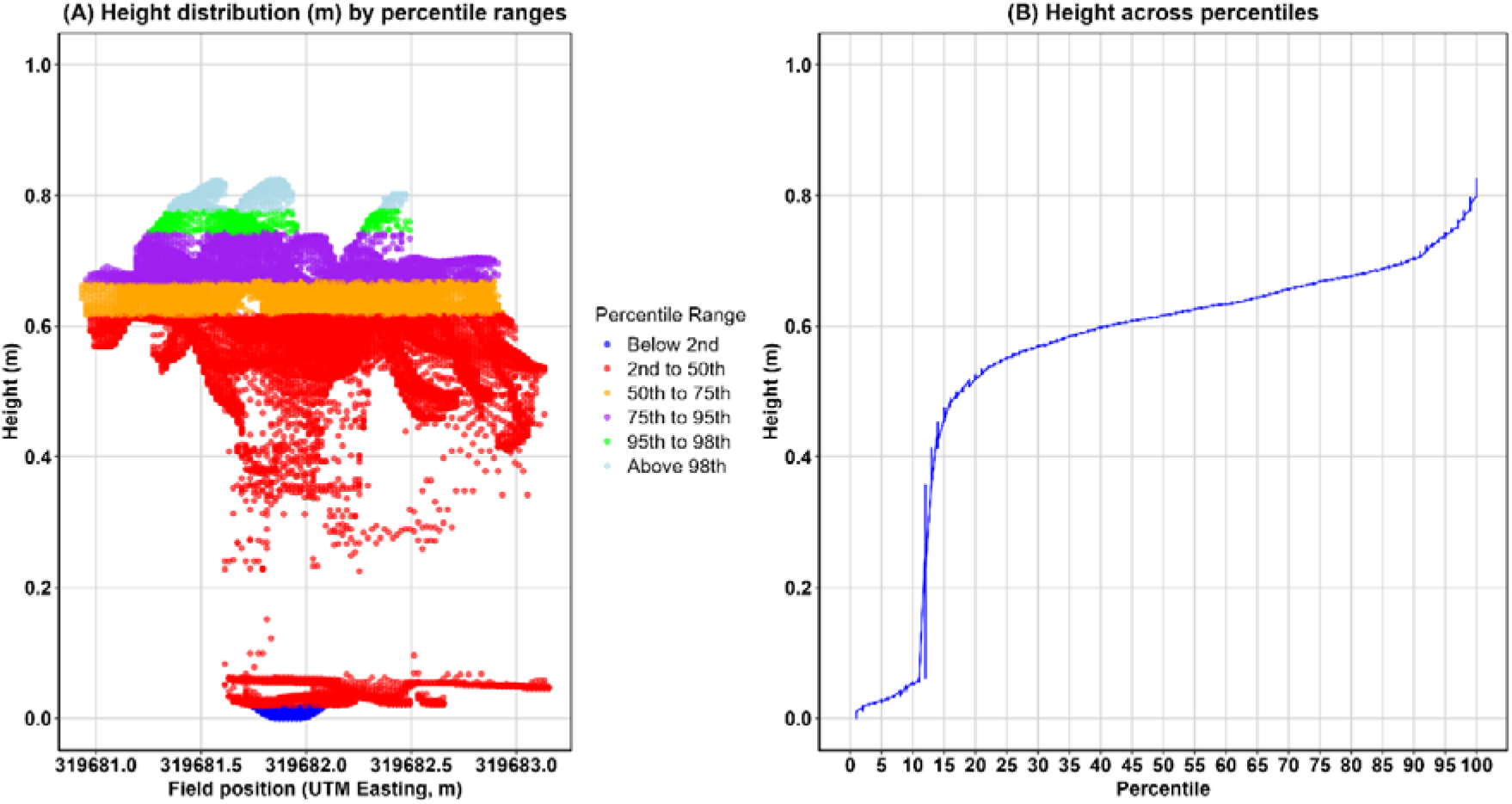
Height data for a representative plot at the flowering stage. (A) Spatial distribution of height measurements along the field position (UTM Easting, m), with points color-coded by percentile ranges. (B) Height values across percentiles (0-100), showing the empirical height–percentile relationship.

This approach maintained all visible ground surfaces in the data, with the height values from the 1^st^ to 99^th^ percentiles representing both ground and plant material, thus capturing the complete vertical profile of the plot.

The general formula for calculating percentile heights is given by:

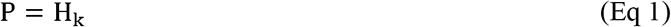

Where:

- P represents the height corresponding to the desired percentile
- H is the set of all height values in the region of interest (e.g., 1 to 99).
- heights are sorted in ascending order: h_1_ < h_2_ < h_3_ < h_4_ … <h_n_
- The index k is calculated as: k = [0.95 * n] (for the 95^th^ percentile height)
- n is the total number of pixels.

The NumPy percentile function calculates the specified percentile by interpolating the sorted height values at the corresponding index. This approach offers a reliable method for determining key height thresholds within the region of interest (Eq 1).

#### 2.3.2 Data cleaning

Data cleaning was performed on the UAV-based photogrammetry data of PHI from the first flight using Tukey’s method for outlier detection. The interquartile range (IQR) was calculated separately for the 75^th^ and 95^th^ percentile heights at flight 1. Based on Tukey’s method, plots outside the lower and upper limits of Q1 - 1.5 × IQR and Q3 + 1.5 × IQR were first identified as potential outliers (Fig S2). After this, the relationship between the 75^th^ and 95^th^ percentile heights was checked visually, because these two percentile heights are expected to have a strong positive relationship in normal plots. The UAV-derived plot images and point cloud data were also inspected to determine whether the flagged plots represented true biological variation or measurement and reconstruction errors. Only plots that were identified as outliers by Tukey’s method and also showed clear deviation from the expected relationship between the 75^th^ and 95^th^ percentile heights were removed. The cleaned dataset was then used for all subsequent analyses (Fig S3).

### 2.4 Differences in percentile height index (DPHI) between two flights

As mentioned above, PHI was extracted from point cloud images obtained during two UAV flights. In non-lodged plots, certain percentiles have been shown to correlate with actual canopy heights.

The left panel (Fig 2A) demonstrates the height distribution when the crop was at its maximum height during the flowering stage (flight 1), with no lodging. The right panel (Fig 2B) shows a change in the height distribution for the same plot after lodging occurred (flight 2). The change in canopy height across all percentiles is due to lodging. Thus, the difference in percentile heights between the two flights shows as a clear indicator of lodging severity.

**Fig 2:**
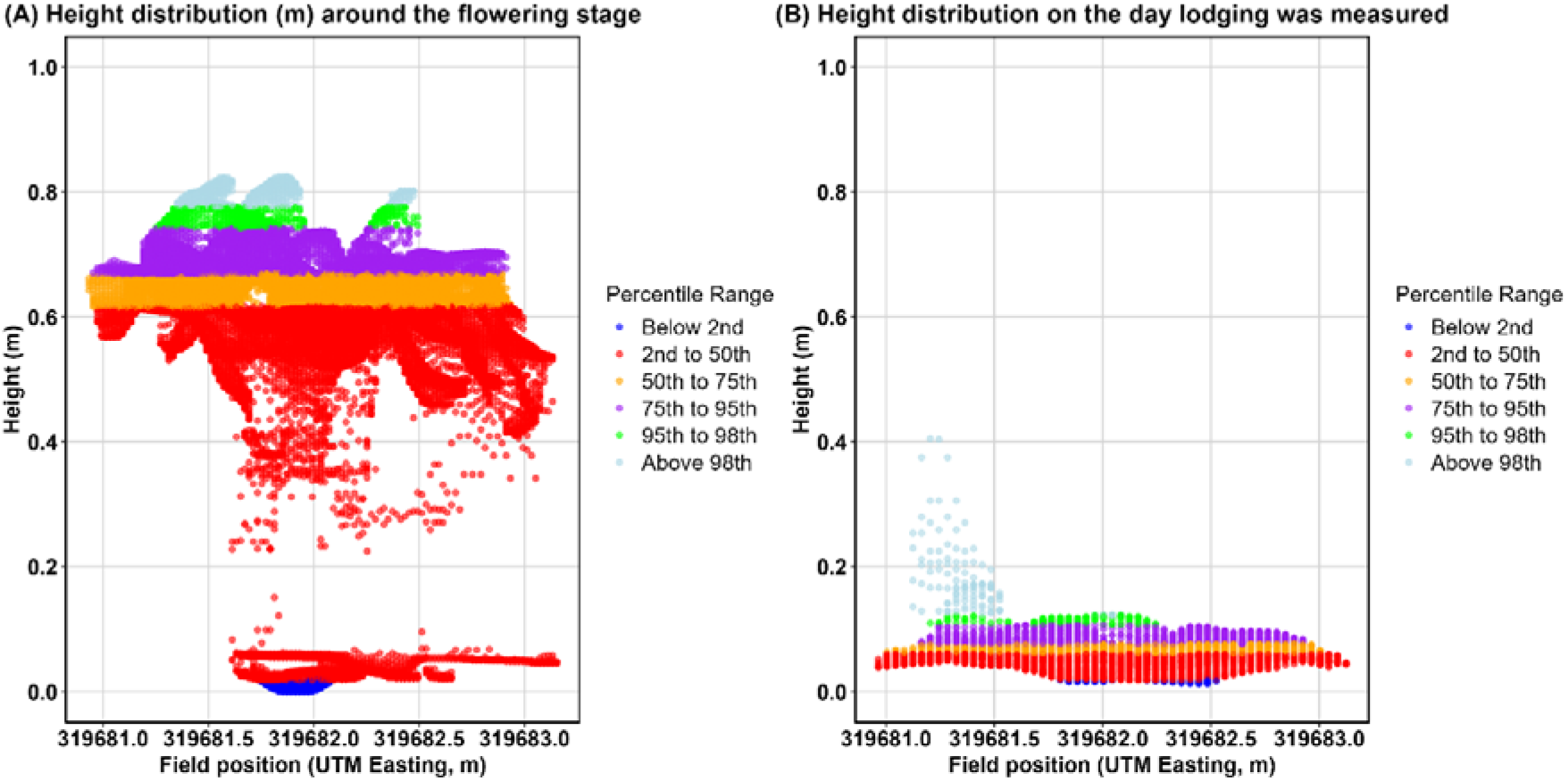
(A) Height measurements for a plot around the flowering stage plotted against field position (UTM Easting, m), with points color-coded by percentile ranges. (B) Height measurements for the same plot plotted against the same field position on the date lodging was assessed (manual lodging severity = 90%). Similar percentile-based height distributions for plots with manual lodging severities of 0%, 60%, and 90% are provided in the supplementary materials (Figs S4-S6).

The differences were calculated using the formula:

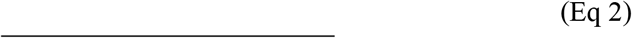

Where:

- is the difference in percentile height index at the 95^th^ percentile between flight 1 and flight 2
- PHI95_Flightt2_ is the height at the 95th percentile in flight 2
- PHI95_Flightt2_ is the height at the 95^th^ percentile in flight 1

We assumed differences in percentile heights between flight 1 and flight 2 are indicators of lodging, with DPHI95-95 quantifying the extent of lodging within the plot. To normalize height variability across plots and improve comparability, the differences were divided by the flight 1 percentile height measurements (Eq 2). This normalization reduced the influence of plant height variability, enabling the evaluation of lodging impact.

### 2.5 Selection of predictors from UAV-based photogrammetry methods

In model construction, selecting efficient predictors is as essential as choosing the algorithm itself. In supervised learning, predictor selection is typically performed before model development to reduce the feature set’s dimensionality, thereby optimizing algorithm performance. In this study, we created several predictors based on the percentile heights between two flights. The variability among predictors was assessed to identify a subset that minimised redundancy while retaining most of the underlying variation. Principal Component Analysis (PCA) was used for this purpose, providing a way to reduce dimensionality within an orthogonal space while preserving most of the original information. PCA highlights the traits that contribute most strongly to variation in the dataset. The analysis was performed using the prcomp function in R (R Core Team, 2025), which applies eigen decomposition to the covariance matrix of standardised variables. The resulting principal components are orthogonal linear combinations of the original predictors, with each component capturing a distinct, uncorrelated portion of the total variance.

By identifying the components that explain the greatest proportion of variation, this approach enabled dimensionality reduction while retaining the most informative predictors. PCA also provided graphical insights into relationships, clustering patterns, and variability among the predictors and the target trait. The overarching goal was to develop a parsimonious model that uses the smallest number of predictors necessary to maximise explanatory power. Reducing the predictor set in this way lowers redundancy, decreases the risk of overfitting, and improves generalisability, resulting in a more robust and interpretable modelling framework.

Pairwise Pearson correlations were then calculated to quantify the associations between manual lodging counts and each selected predictor.

### 2.6 Prediction approaches to ensure model robustness

To assess model robustness, cross-validation was performed and the distribution of lodging phenotypes in the training data was examined to ensure predictions were not driven by a narrow or biased training range. These methods were designed to ensure a thorough assessment of model performance across varying data conditions and distributions.

#### 2.6.1 Cross-validation approach

A cross-validation approach was utilized to assess the accuracy of predictive models for lodging. The process involved utilizing the total available data from multiple sites and trial series. The complete dataset of all plots from both sites was randomly split into training and testing sets (Fig S7).

The training set comprised 80% of the data, which was used to train statistical models capable of predicting lodging. The remaining 20% of the data was allocated to the testing set (Fig S7). To obtain reliable distributions of cross-validated accuracies, the entire process was replicated for 100 iterations (Fig S7). Each iteration involved a new random partitioning of the data, ensuring that the results were not biased by any specific data split. This iterative approach provided a robust estimate of the model’s predictive performance across different subsets of the data.

#### 2.6.2 Evaluation of distribution of phenotypes in the training set

The research utilized two distinct approaches for creating training sets from the complete dataset of 2,675 plots (Table S1). The first approach involved stratified sampling to ensure equal representation across different lodging severity levels, while the second employed random sampling, which maintains the natural distribution of lodging severity in the dataset utilising the manual lodging counts (Fig S8). To implement these approaches, the plots from all sites were first classified into 11 groups based on observed manual lodging count, where counts represented the percentage of lodged plants (Table S1). For the random sampling approach (Approach 1), a training set comprising 550 samples was randomly selected from all severity groups while preserving the natural distribution of manual lodging counts (Fig S8). The distribution of plots across these groups is illustrated in Supplementary Table (1) . For the stratified sampling approach (Approach 2), we randomly selected 50 samples from each severity group to create a balanced dataset of 550 plots, minimizing potential training biases (Fig S8). An independent test set of 550 randomly selected plots was used for model validation. This methodology was repeated over 100 iterations to evaluate model performance under both balanced and imbalanced training conditions, providing insight into the model’s robustness across different data distributions.

### 2.7. Statistical and mathematical learning approaches

To develop a model to predict lodging at the plot level using the UAV-based photogrammetry method, a range of parametric and non-parametric statistical learning as well as a mathematically driven ensemble approach to predict lodging were evaluated (Hastie et al. 2009). The manual lodging counts and selected predictors were used as inputs for constructing these models. Models were implemented in R (R Core Team, 2025), and with the caret package (Kuhn 2008), which enabled the development of models that optimize the prediction of manual lodging counts across various conditions. In each model, a seed was set for reproducibility, and 1000 permutations of fivefold cross-validation were conducted on the training set using a tuning grid of model-specific hyperparameters to select the optimal hyperparameters (Table 2) based on the lowest root mean square error (RMSE) achieved. The mean performance of each model was calculated by averaging the performance across individual folds and repeats, thereby minimizing prediction bias introduced by random sampling in this relatively small dataset.

**Table 2:**
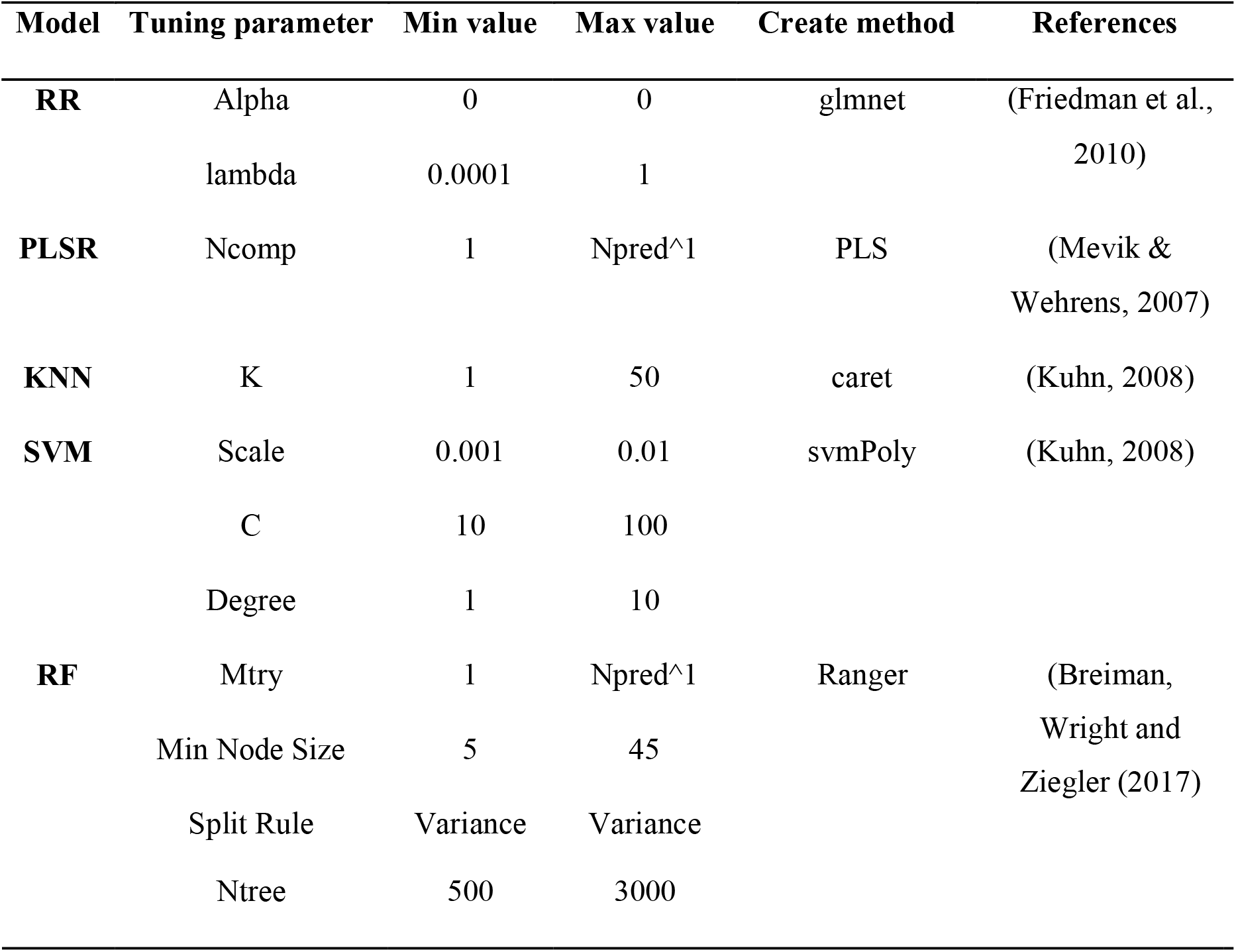
Overview of hyperparameters used in cross-validation for each model; Npred: Number of predictors; Mtry: Number of features considered at each split; Ncomp: Number of components; Npred: Number of predictors; Ntree: Number of trees.

#### 2.7.1 Parametric models

Parametric models assume consistent responses across conditions, including MLR, RR, and Partial Least Squares regression (PLS).

MLR analyses relationships between lodging (dependent variable) and predictor variables, assuming linear and additive effects, and uses ordinary least squares (OLS) for simplicity and optimality.

The general MLR model is:

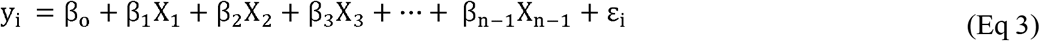

Components:

- y_i_ represents lodging on plot *i*
- β_0_ is the intercept
- β_n_ the regression coefficients
- X_n_ are the DPHI predictors
- ε_i_ the residual error
- (ε_i_ ∼ N 0,σ^2^) is the residual error on plot i, normally distributed with a mean zero and a constant variance.

Regression coefficients (Eq 3) quantify the linear impact of explanatory variables on lodging, providing insights into how predictors influence lodging. RR addresses multicollinearity by adding bias to regression estimates, reducing variance, and improving model stability and generalizability by shrinking coefficients (Hoerl & Kennard 1970). PLS estimate predictors and response variables onto uncorrelated latent variables, simplifying models, capturing key data relationships, and enhancing predictive accuracy, especially with high collinearity. Cross-validation determines the optimal number of components, ensuring robust models (Geladi & Kowalski 1986).

#### 2.7.2 Non-parametric models

Non-parametric models do not rely on assumptions regarding the underlying distributions of the model components. Instead, the model parameters are learned through an iterative training process. The applied methodology included KNN, SVR, and RF selected from various machine-learning models.

The KNN regression method assumes that similar samples are located near the K closest samples in the feature space. The value of K is a hyperparameter that must be tuned for each dataset. This study examined values of K from 1 to 50 in increments of five (e.g., K = 1, 5, 10, 15, 20, 25, 30, 35, 40, 45, 50) (Table 2). The optimal value of K was derived from cross-validation, minimizing the residual mean square error.

The SVR model with a radial basis function (RBF) kernel is a machine-learning technique designed to capture non-linear relationships by mapping the input data into a higher-dimensional feature space. In this space, the algorithm learns a regression function that fits within an ε-insensitive margin, where only data points lying outside this tube become support vectors. These support vectors determine the shape of the regression function. By maximising the margin around the function while minimising prediction error, the model achieves a smooth, generalisable fit capable of modelling complex non-linear patterns.

RF is an ensemble learning method that comprises a collection of decision trees (Breiman 2001). Each decision tree follows a hierarchical tree-like structure made up of nodes and edges. The structure begins at the root node, which serves as the starting point. This root node splits into two edges, leading to decision nodes and eventually to terminal nodes, or leaves, which represent the final predicted outcome. A decision node acts as an intermediate step where a condition is evaluated based on the values of a specific feature in the dataset. If the condition is satisfied, the algorithm moves along the left edge to the next node; otherwise, it moves to the right. Each tree in the RF is trained on a unique sub-training set sampled with replacement, ensuring diversity among the trees. The parent-child relationship is maintained within the tree, with the root node acting as the parent node to its subsequent branches, including sub-trees, intermediate nodes, and terminal leaves. The final RF prediction is generated by aggregating predictions from all individual trees, effectively reducing prediction noise and enhancing stability.

#### 2.7.3 Ensemble naïve average

To enhance prediction accuracy and improve generalization, we implemented an ensemble approach which potentially leverages the strengths of each of the algorithms we used (Tomura et al. 2025).

In this approach, independent predictions were generated using six distinct models: MLR, RR, PLS, KNN, SVM radial, and RF. These predictions were then averaged to produce a naïve average (Fig 3). The term naïve was used to indicate the fact that no attempt was made to weight the predictions based on the contributions of the individual prediction models. Each model independently predicted phenotypes using the UAV-enabled photogrammetric method. The outputs from these models were assembled into a predicted phenotype matrix (Fig 3), where each row contained the predictions of the individual models for a specific target. The ensemble naïve averaging method was then applied by averaging the predictions from all models, giving equal weight to each. This approach leverages the complementary strengths of the models to produce robust and reliable predictions. These predictions were evaluated using performance metrics to assess prediction accuracy.

**Fig 3:**
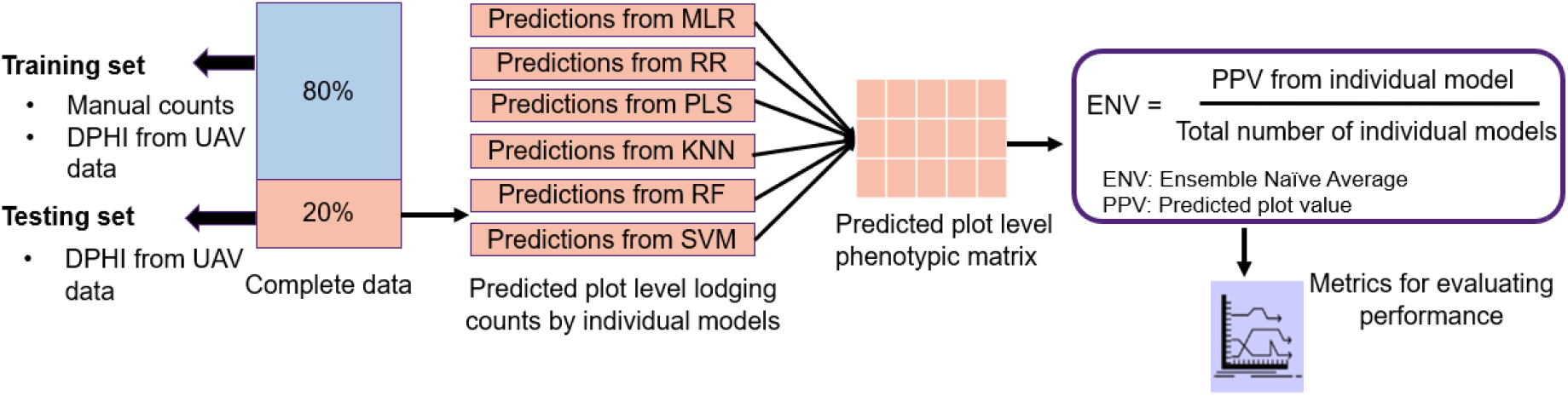
Schematic illustration of ensemble naïve averaging using UAV data and multiple models from cross-validation approach [Image idea: (Tomura et al. 2025)].

#### 2.7.4 Metrics for evaluating model accuracy

In this study, four key metrics were employed to evaluate the predictive performance of the models: residual squares, RMSE, Pearson correlation coefficient (PCC), and coefficient of determination (R²) derived between observed and predicted plot values. A statistical threshold was also applied to identify high-residual plots.

Residual squares were calculated as the difference between observed and predicted lodging values, and the squared residuals were computed to evaluate prediction accuracy

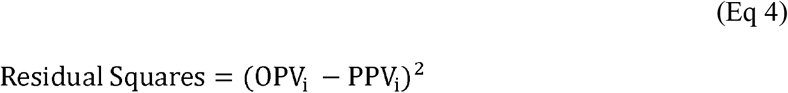

Where:

- OPV_i_ and PPV_i_ represent the observed and predicted plot values, respectively,

The RMSE metric, defined as the square root of the average squared differences between the observed and predicted values, provides a quantification of the distance between observed and predicted values (Eq 5).

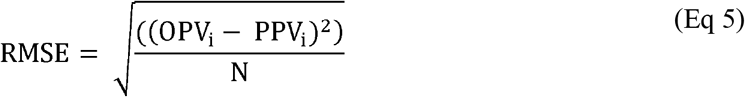

Where:

- N is the total number of plots in the testing set

Lower residual squares and RMSE values indicate smaller differences between observed and predicted values, corresponding to higher predictive accuracy.

The PCC was calculated to measure the strength and direction of relationship between the observed and predicted plot values. The PCC value ranges from -1 to 1, with values closer to 1 signifying a stronger positive correlation, thus reflecting higher predictive accuracy.

The R² evaluates the proportion of variance in the dependent variable explained by the predictive model. R² ranges from 0 to 1, with values closer to 1 indicating that the model accounts for a larger proportion of the variability in the observed data. An R² value of 1 represents perfect prediction, while 0 indicates that the model explains no variability.

A stringent RMSE-based cutoff was used to identify plots with extreme prediction errors. First, RMSE was calculated for each of the 100 iterations and the mean RMSE was obtained.

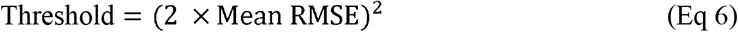

Plots with residual squares above this threshold were further investigated (Eq 6). Plots were classified as high residual (poorly predicted) when their residual squares exceeded the threshold.

#### 2.7.5. Iterative refinement and validation

To investigate sources of prediction error, plots with large residuals were identified based on model prediction errors. UAV imagery for each of these plots was visually inspected and compared against manual lodging counts. Plots were classified as having either (i) incorrect manual measurements; (ii) poor crop establishment, defined as establishment scores of 4 to 6. To assess the impact of these error sources on model performance, plots identified as incorrectly measured and poorly established were removed individually and jointly from the dataset. Model performance was then re-evaluated by recalculating RMSE using the original trained models. In addition, all models were retrained using a modified training dataset that excluded plots with confirmed manual measurement errors and poor establishment. Based on this procedure, plots with poor establishment (scores 4-6) and plots with confirmed manual measurement errors were excluded from subsequent analyses.

## 3. RESULTS

### 3.1 Distribution of manual lodging counts (%) across environments

The distribution of manual lodging counts differed substantially between Jandowae and Pirrinuan. At Jandowae, approximately 500 plots (the highest frequency) showed lodging counts of 0-5%, followed by about 320 plots at 5-10%, with frequencies rapidly declining thereafter (Fig 4).

**Fig 4:**
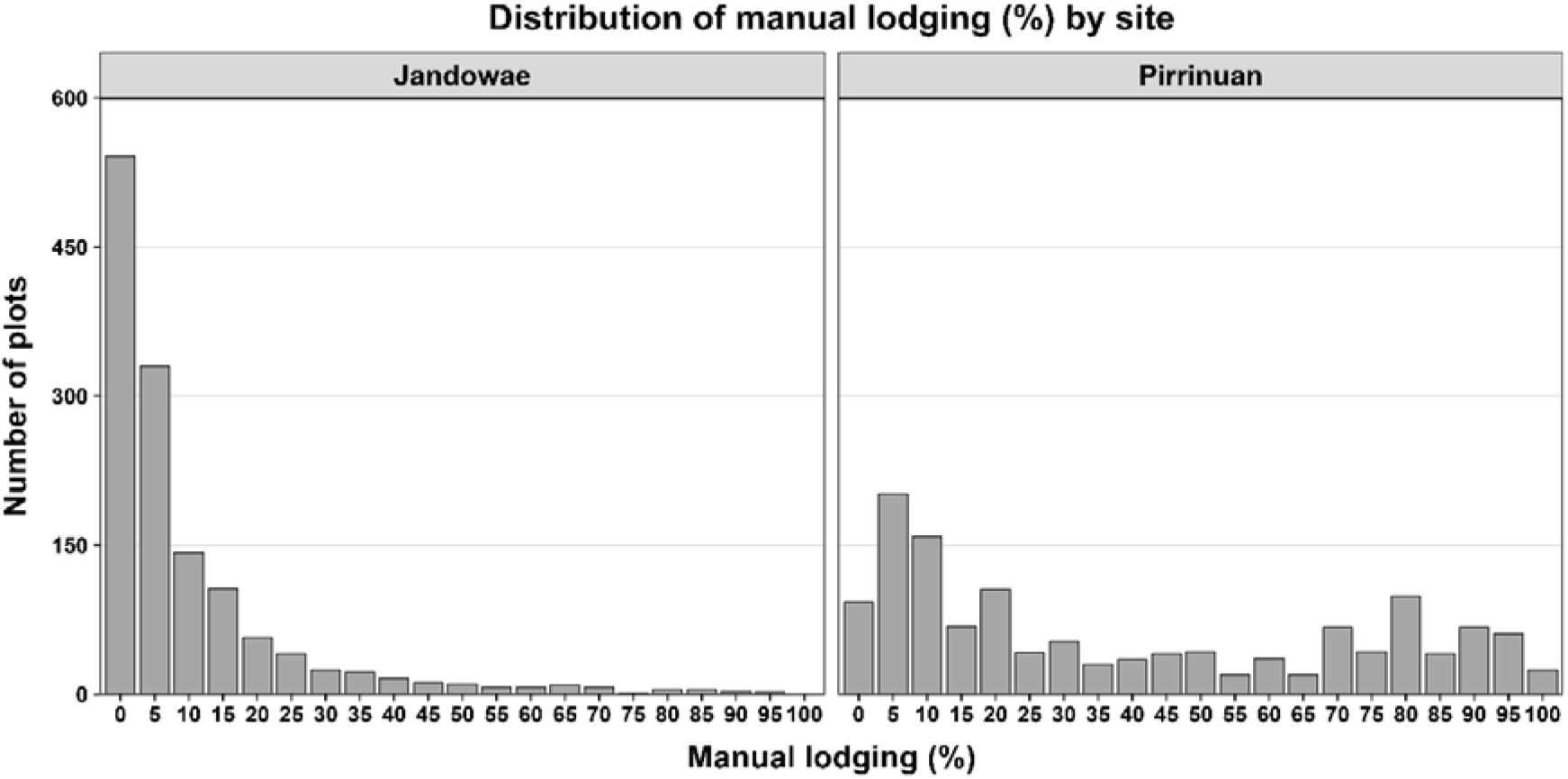
Distribution of manual lodging percentages across plots at Jandowae and Pirrinuan.

At Pirrinuan, the phenotypic distribution showed two main peaks: one with approximately 200 plots showing 10-15% lodging, and another with about 100 plots in the 80-90% range (Fig 4). The remaining manual lodging counts at Pirrinuan were more evenly distributed, with frequencies generally ranging from 20 to 50 plots (Fig 4).

### 3.2 Exploratory assessment and selection of UAV-derived canopy-height predictors

PCA was used as an exploratory analysis to visualise relationships and redundancy among the UAV-derived photogrammetric variables. In the initial analysis PC1 and PC2 explaining 74.2% and 18.1% of the total variance (Fig 5A).

**Fig 5:**
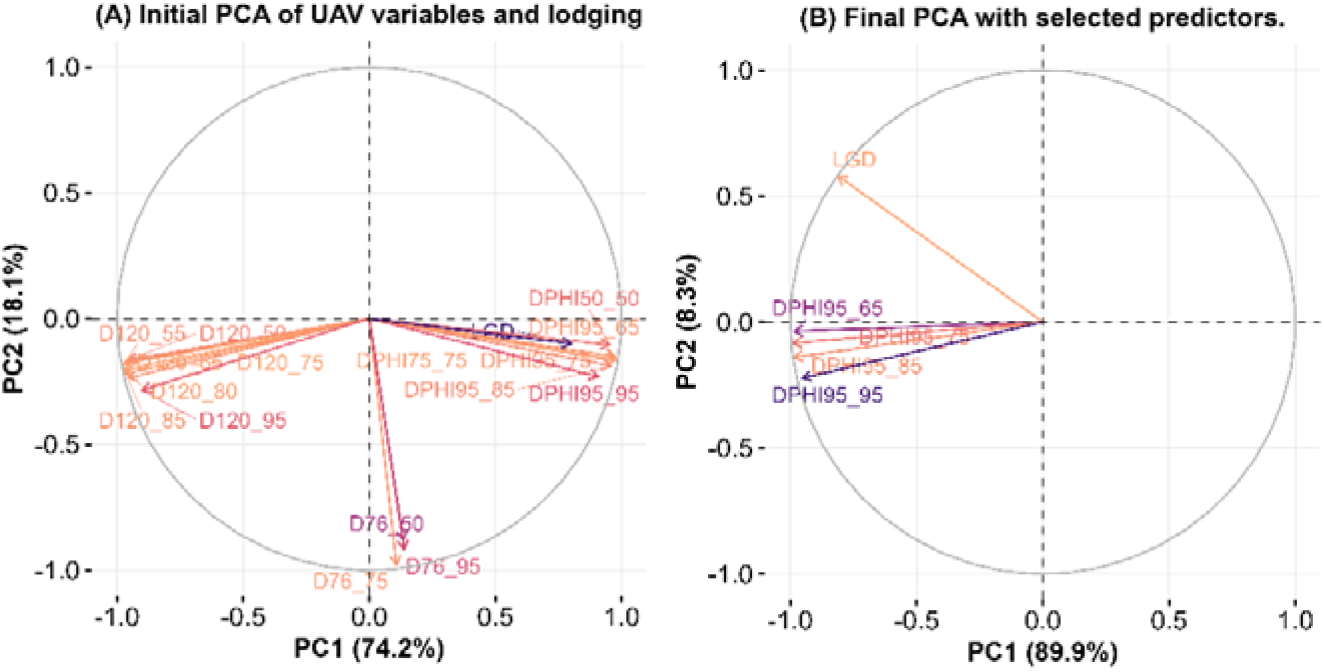
PCA biplots for predictor selection. (A) Initial PCA incorporating all UAV-enabled photogrammetric variables, showing the relationships among multiple predictors. (B) Final PCA highlighting the selected variables used in model development for lodging prediction. Here, D120_75 represents a predictor derived from flight 2, based on the 75^th^ percentile height after manual lodging counts were taken, while D76_75 represents a predictor derived from flight 1, based on the 75^th^ percentile height around the flowering stage.

In the initial PCA variables (Fig 5A) showed distinct clustering patterns, with some variables exhibiting high correlations as indicated by similar vector directions (e.g. DPHI95 measurements at different height thresholds). The D120 series variables formed a separate cluster, suggesting they capture different aspects of canopy compared to the DPHI95 series. Based on this analysis, we selected DPHI95_65, DPHI95_75, DPHI95_85, and DPHI95_95 as the final predictors for predicting lodging (LGD, the target variable). This selection was guided by the strong loadings of these variables on the principal components (indicated by vector length). The final PCA (Fig 5B) shows that these selected predictor variables explain 89.9% of the variance in PC1 and 8.9% in PC2, indicating that the selected subset effectively captures the majority of the variation in the dataset while reducing dimensionality.

Pairwise Pearson correlations were then used to quantify these relationships (Fig 6). Manual lodging was positively correlated with DPHI95_65, DPHI95_75, DPHI95_85, and DPHI95_95, with correlation coefficients of 0.769, 0.751, 0.721, and 0.656 (Fig 6). The correlation with manual lodging gradually decreased from the 65^th^ to the 95^th^ percentile. The DPHI95 variables were also highly correlated with each other, with correlation coefficients ranging from 0.916 to 0.993 (Fig 6). The selected DPHI variables remained highly correlated because they represented different percentile levels of the same canopy-height distribution. However, the correlations were below 1.0, and their positions in the PCA space indicated that the percentile levels were not completely identical in the information they represented.

**Fig 6:**
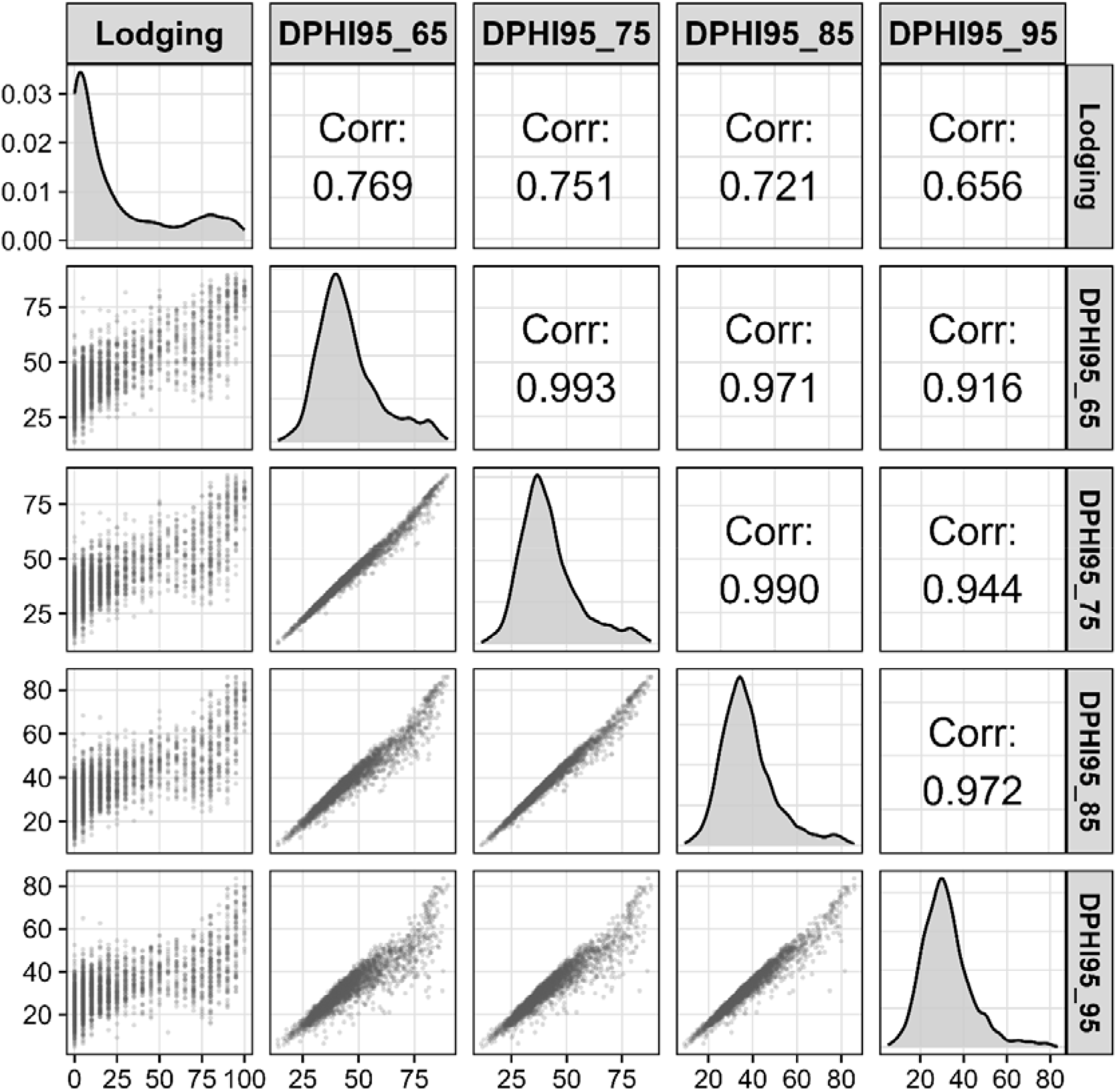
Pairwise plot matrix showing the distributions, scatterplots, and Pearson correlations among manual lodging counts and the four DPHI95 variables.

These results showed that the UAV-derived photogrammetric height-change variables were moderately to strongly associated with manual lodging counts. Although all four DPHI variables were associated with manual lodging, their relationships with lodging were not necessarily strictly linear and the predictors contained substantial shared information. We therefore evaluated both parametric and non-parametric modelling approaches, together with an ensemble approach, for plot-level lodging prediction.

### 3.3 Predicting sorghum lodging from UAV-enabled photogrammetric methods

This study evaluated multiple statistical models for predicting crop lodging using a cross-validation approach that integrated manual lodging counts with UAV-enabled photogrammetric metrics across different sites. The comparative analysis revealed distinct performance patterns among parametric, non-parametric models and ensemble naïve average (Fig 7).

**Fig 7:**
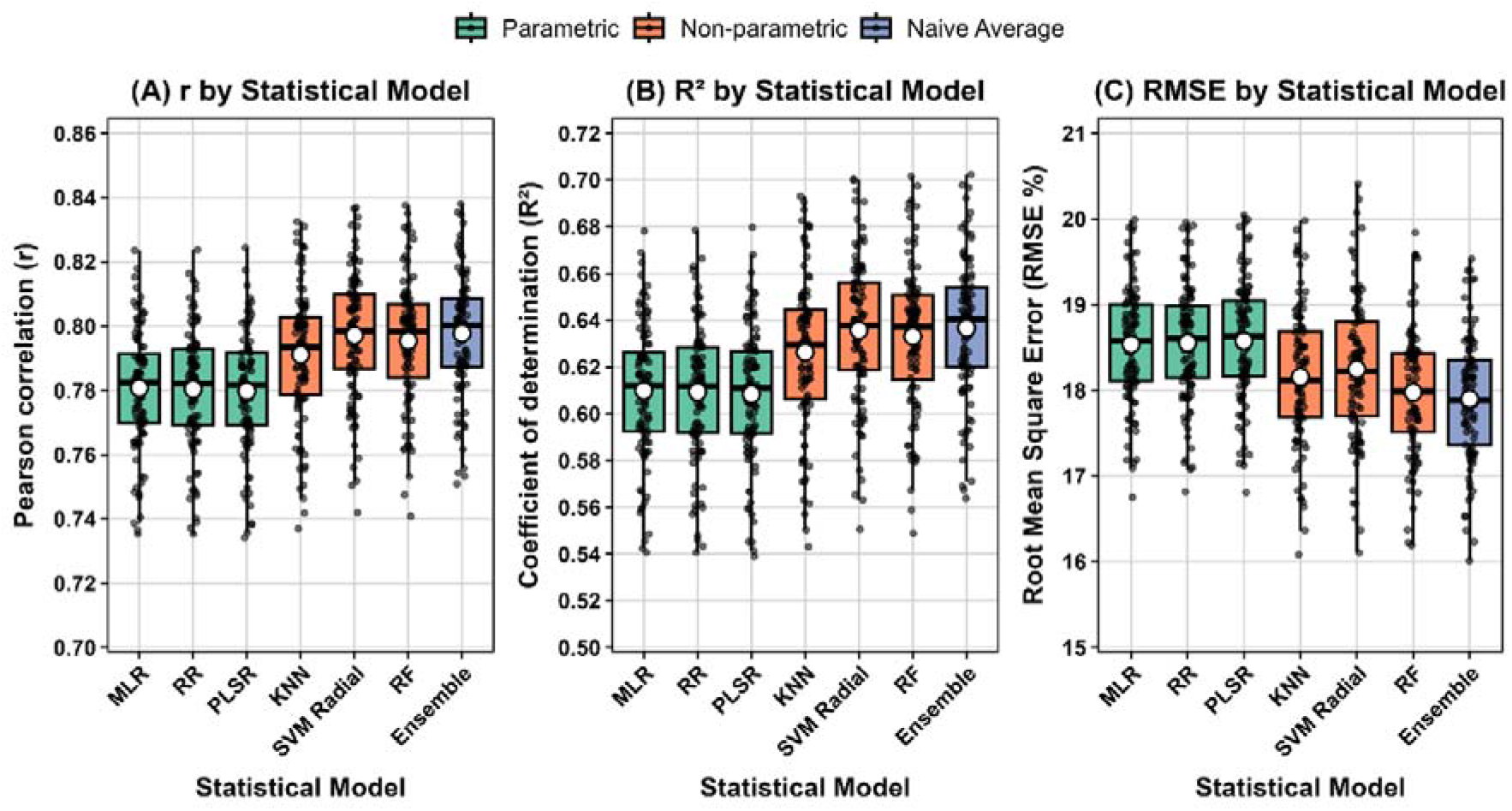
Comparative cross-validation performance of statistical models evaluated using r, R², and RMSE. Black points show the distribution of model performance across 100 iterations, and red points represent the mean values.

The parametric models (MLR, PLSR, and RR) showed consistent performance patterns, with slightly lower overall r and R² and higher RMSE than the non-parametric models. MLR achieved marginally better error metrics (RMSE = 18.54 ± 0.68) compared to RR (18.55 ± 0.68), while PLSR showed slightly higher variability (18.58 ± 0.68). Despite minimal differences in error metrics, each parametric model extracted slightly different patterns while maintaining similar explanatory power (R² = 0.61) (Fig 7 and Table S2). Among the non-parametric models, SVM Radial and RF showed strong performance, with mean correlations of 0.80 ± 0.02, respectively. However, the SVM Radial model exhibited higher prediction variability (RMSE = 18.25 ± 0.84) compared to RF (RMSE = 17.98 ± 0.73). KNN model demonstrated intermediate performance (r = 0.79 ± 0.02, r² = 0.63 ± 0.03, RMSE = 18.16 ± 0.79) (Fig 7 and Table S2).

The ensemble naïve average method demonstrated slightly superior performance across all evaluation metrics, achieving a r = 0.8 ± 0.019 and R^2^ = 0.64 ± 0.03, while maintaining the lowest RMSE = 17.90 ± 0.73 (Fig 7 and Table S2). Its compact error distribution and high correlation values suggest superior stability and predictive capability compared to individual models. These findings show that although the naïve ensemble delivers the most balanced and consistently strong performance for lodging prediction, both parametric and non parametric models remain viable options depending on the desired balance between accuracy, interpretability, and computational efficiency.

### 3.4 Prediction accuracy and identification of poorly predicted plots using Ensemble naïve average model

The developed ensemble model was used to investigate the impact of lodging severity on prediction accuracy and to identify sources of error contributing to poorly predicted plots across the entire dataset.

#### 3.4.1 Prediction accuracy across lodging severity categories

The prediction accuracy of the ensemble naïve average model varied substantially across lodging severity categories. A threshold of 1282 for residual squares, calculated using equation (6), was used to classify plots as “poorly predicted”.

The ensemble model demonstrated high prediction accuracy in the no lodging category (0%), with 98.9% of predictions falling below the threshold (Fig 8 and Table S3). Performance remained consistently strong for low lodging categories (5-30%), with 97.4-98.7% of predictions below the threshold (Table S3), although slightly increased variability was observed, with a few plots reaching residual squares up to 6,000, particularly in the 5-10% category (Fig 8). Through moderate lodging ranges (35-50%), prediction accuracy remained relatively stable, with 92.4-100% of predictions falling below the threshold (Fig 8 and Table S3). However, a notable decline in prediction accuracy occurred in higher lodging categories (55-80%). In the 55-60% category, 76.7% of predictions remained below the threshold, decreasing to 60.8% and 56.3% in the 65-70% and 75-80% categories, respectively (Fig 8 and Table S3).

**Fig 8:**
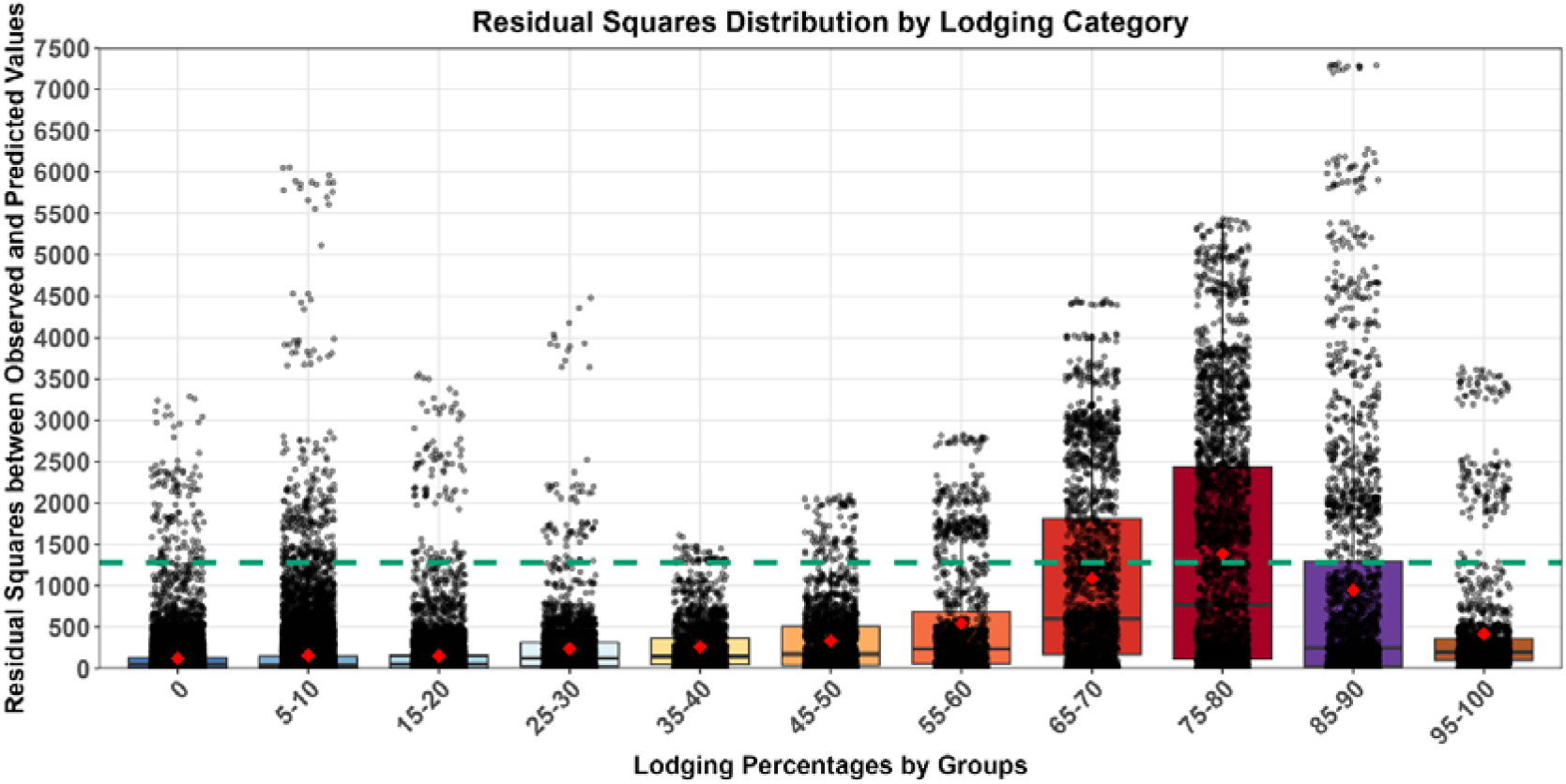
Distribution of residual squares across lodging categories from the ensemble naïve-average model, based on 100 cross-validation iterations. The green line marks the threshold (1282) used to identify poorly predicted plots.

Interestingly, prediction accuracy recovered in the most severe lodging categories (85-100%). The 85-90% category showed 78.6% of predictions below the threshold, while the highest category (95-100%) demonstrated improved performance with 95.8% of predictions below the threshold (Fig 8 and Table S3). Residual-based performance tended to decline through some intermediate-to-high lodging categories and improved again in the most severe category. However, the number of plots differed substantially among lodging classes. Overall, 93.2% of predictions (2,495 out of 2,675 plots) fell within the threshold, while 6.8% (180 plots) were identified as poorly predicted.

#### 3.4.2 Investigation of poorly predicted plots

Among the 180 poorly predicted plots identified from the complete dataset of 2,675 plots, we conducted a detailed visual inspection using plot-level images extracted from the UAV imagery. For each poorly predicted plot, the image covering the two-row plot was compared with both the manually measured lodging count and the UAV-predicted lodging count. Examples of this visual inspection are provided in the Supplementary Information (Figs. S9-S14). This assessment identified two main factors associated with poor prediction performance (Table S4).

The first factor was potential transcription or recording errors in the manual lodging data. This was observed in 82 plots, representing 3.1% of the complete dataset (2,675 plots). For these plots, visual inspection indicated that the recorded manual lodging counts did not correspond with the lodging condition visible in the single-plot images extracted from the UAV imagery (Figs. S9-S13). These discrepancies were therefore considered likely to reflect transcription, recording, or plot-assignment errors rather than errors in the plant-counting procedure itself.

The second factor was poor establishment, which was observed in 29 plots, representing 1.1% of the complete dataset (2,675 plots) and 16.1% of the 180 poorly predicted plots (Table S4). Poor establishment was defined by establishment ratings of 4, 5 or 6 (Fig S14). Plots with poor establishment occurred more frequently among the poorly predicted plots than expected based on their proportion in the complete dataset. In breeding trials, grain yield, plant height, flowering and other trait measurements from poorly established plots may not represent the genetic performance of the tested material.

For the remaining 69 poorly predicted plots, no obvious cause of error was identified in 59 plots. In five plots, the poor predictions were associated with errors in the UAV-based lodging estimates, while plot-level images could not be extracted for visual inspection for the remaining five plots.

After identifying the main sources of error among the poorly predicted plots, we excluded 82 plots with potential transcription or recording errors in the manual lodging data and 162 plots with poor establishment ratings of 4-6 from the complete dataset of 2,675 plots. The parametric, non-parametric and ensemble models were then retrained using the remaining plots, and their prediction performance was evaluated (Fig S15 and Table S5). For the selected ensemble naive-average model, Pearson’s correlation coefficient ranged from 0.82 to 0.89, while RMSE ranged from 13.00 to 15.96 percentage points across the prediction scenarios (Table S5).

### 3.5 Impact of transcription errors and poor establishment on model performance

Prediction accuracy differed depending on the source of error present in the data (Fig 9; Table S6). The original ensemble model trained using the complete dataset the identified problematic plots, achieved a mean RMSE of 17.90 (± 0.73) percentage points across 100 iterations, with values ranging from 16.01% to 19.54 percentage points.

**Fig 9:**
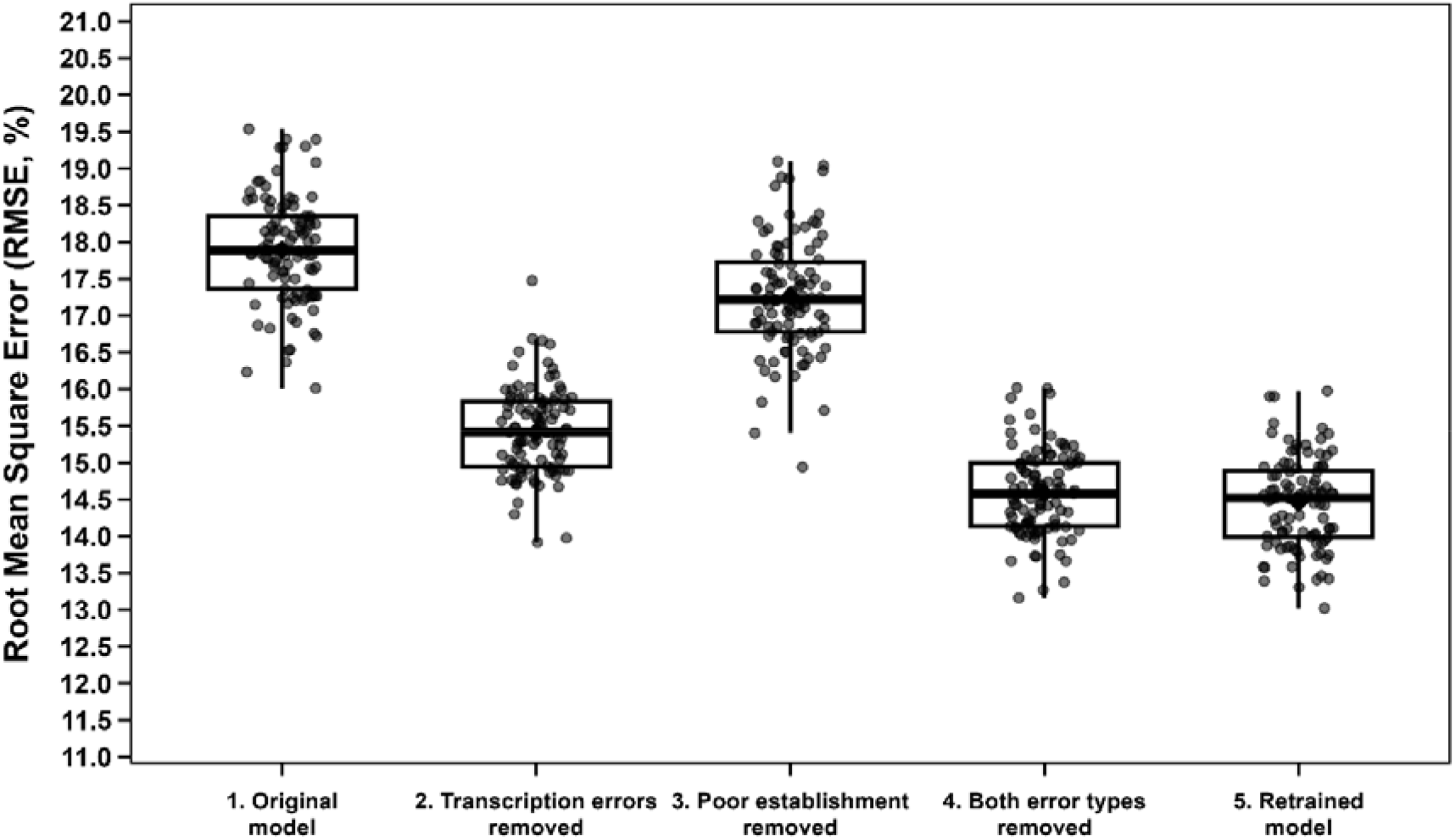
Comparison of ensemble model prediction accuracy (RMSE) across five approaches: (1) original model; (2) original model with transcription-error plots removed; (3) original model with poor-establishment plots removed; (4) original model with both transcription-error and poor-establishment plots removed; and (5) retrained model after removing both error types.

When plots affected by potential transcription or recording errors in the manual lodging data were removed from the evaluation dataset, mean RMSE decreased substantially to 15.42 (± 0.60) percentage points, with values ranging from 13.92 to 17.48 percentage points. In comparison, removing plots with poor establishment alone resulted in a smaller improvement, with mean RMSE decreasing to 17.27 (± 0.77) percentage points and ranging from 14.94 to 19.09 percentage points. These results indicate that potential transcription or recording errors in the manual lodging data contributed more strongly to prediction error than poor establishment.

When plots affected by both potential transcription or recording errors and poor establishment were removed, prediction accuracy improved further, with mean RMSE decreasing to 14.59 (± 0.59) percentage points and values ranging from 13.16 to 16.02 percentage points. These first four approaches quantified how much of the observed prediction error was associated with the two identified data-quality issues, without changing the model used to generate the predictions.

To determine whether the presence of these problematic plots during model training influenced model performance, the ensemble model was then completely retrained after removing both error types from the training data. The retrained model achieved a mean RMSE of 14.46 (± 0.62) percentage points, which was very similar to the RMSE of 14.59 percentage points obtained when the problematic plots were simply removed from the evaluation dataset without retraining. This similarity indicates that including a relatively small number of plots affected by transcription or recording errors and poor establishment during initial model training had little influence on the fitted ensemble model. Therefore, the ensemble model appeared relatively consistent to these data-quality issues.

After removal of the identified problematic plots and model retraining, the prediction performance of the individual statistical learning models also improved (Fig S10; Table S5). The performance advantage of the ensemble model over the best-performing individual models became smaller after data cleaning, with the ensemble naïve-average model performing similarly to the SVM radial and RF models. Nevertheless, the ensemble approach was retained for subsequent analyses because it provided consistently strong prediction performance across the different evaluation scenarios.

### 3.6 Spatial comparison of manual lodging counts and UAV-predicted lodging counts

The spatial distributions of manual lodging counts, UAV-predicted lodging counts, and plot-level prediction errors were compared across the four-trial series (Fig. 10 and Figs. S16-S18). The manual and UAV-predicted lodging maps showed broadly consistent spatial patterns across the field layouts, with areas of higher manual lodging generally corresponding to areas of higher UAV-predicted lodging. However, differences between manual and predicted lodging values were evident for some individual plots.

**Fig 10:**
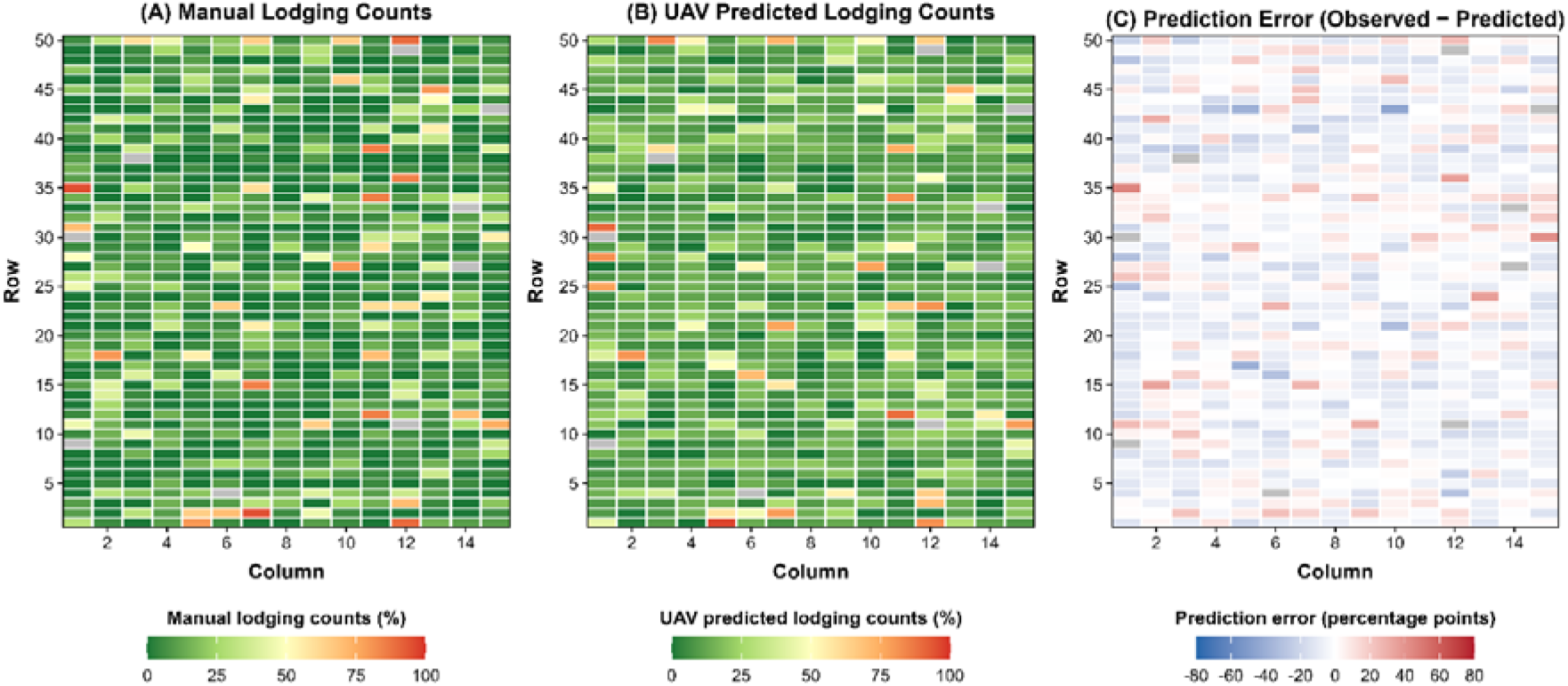
Spatial comparison of observed and UAV-predicted lodging in the AYTF trial at Jandowae. (A) Manually observed lodging (%), (B) UAV-predicted lodging (%), and (C) plot-level prediction error calculated as predicted minus observed lodging (%). Positive values indicate overestimation, while negative values indicate underestimation. Each tile represents an individual plot in the field layout.

The prediction-error maps showed both positive and negative errors across all trial-environment combinations. Positive values indicated overestimation of lodging by the UAV-based model, whereas negative values indicated underestimation. Most prediction errors were distributed across the field layouts, while some individual plots showed relatively large deviations between manual and UAV-predicted lodging. The error maps also allowed visual assessment of whether these larger prediction errors occurred as isolated plots or showed localized spatial patterns within the trials. These spatial comparisons complemented the overall model-performance statistics (r, R², and RMSE) by illustrating the magnitude, direction, and spatial distribution of prediction errors at the individual-plot level.

### 3.7 Site-specific prediction performance and cross-site validation

Fig 11 and Table S7 show the distribution of squared residuals across lodging severity percentages for Jandowae and Pirrinuan using predictions from the retrained ensemble naïve-average model. A threshold of 833 for residual squares, calculated using equation (6), was used to determine the percentage of plots that were poorly predicted.

**Fig 11:**
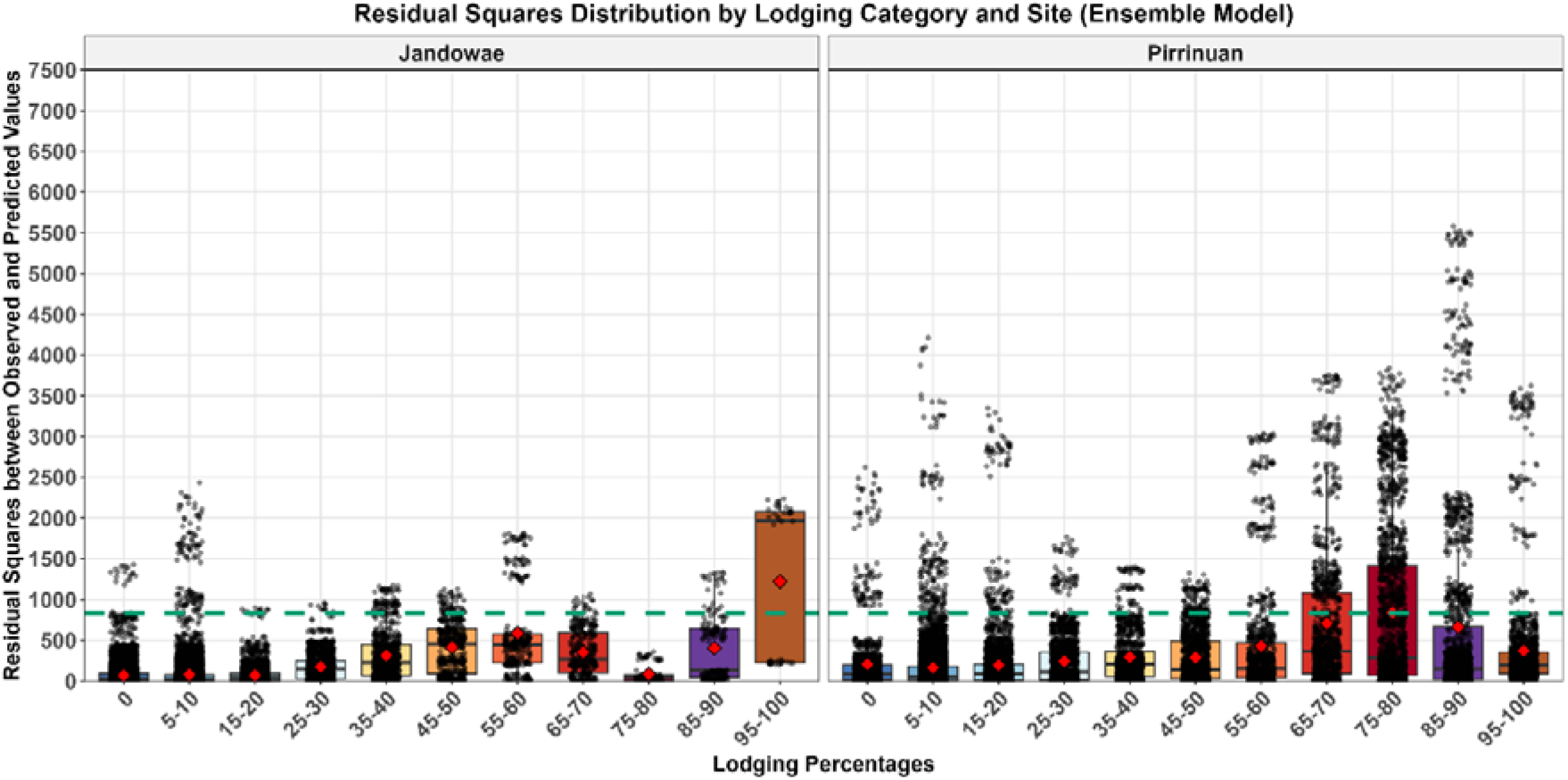
Distribution of squared residuals from the retrained ensemble naïve-average model across lodging severity categories at Jandowae and Pirrinuan. The boxplots represent the residual squares across various lodging percentages, with the red points representing the mean residual squares from 100 iterations. The green line marks the threshold (833) used to identify poorly predicted plots.

For no lodging (0%) and low lodging (5-30%), prediction performance was generally strong at both sites (Fig 11). At Jandowae, 98.7-100% of predictions had squared residuals below the threshold, while at Pirrinuan, 92.6-96.9% were below the threshold (Table S7). For moderate lodging percentages (35-60%), the proportion of predictions below the threshold ranged from 78.6 to 89.5% at Jandowae and from 85.1 to 94.8% at Pirrinuan, indicating generally comparable prediction performance across the two sites.

At higher lodging severity (65-80%), the residual squares revealed divergent patterns between sites. Jandowae maintained strong prediction performance with 93.8-100% of predictions below the threshold (Table S7). For the very high lodging categories (85–100%), prediction performance was more variable, with 50.0-87.5% of predictions below the threshold at Jandowae and 76.9-92.3% at Pirrinuan. These results indicate that prediction error varied among lodging severity classes and between sites, particularly at higher lodging levels. Some of this variability may also reflect differences in the number and distribution of plots represented within individual lodging categories at each site.

Cross-site validation was then used to evaluate the ability of the ensemble model to transfer between environments (Fig. S19). When the model was trained using Pirrinuan data and used to predict Jandowae, prediction performance was r = 0.77, R² = 0.60, and RMSE = 11.58 percentage points. In the reciprocal analysis, training on Jandowae and predicting Pirrinuan resulted in r = 0.83, R² = 0.69, and RMSE = 20.92 percentage points (Fig S19). Thus, the model retained moderate-to-strong relationship between observed and predicted lodging across both cross-site validation directions, although absolute prediction error differed between sites. The higher RMSE when predicting Pirrinuan indicates that prediction transferability was influenced by site-specific differences in the distribution and severity of lodging, despite the relatively strong correlations observed in both directions.

### 3.8 Effect of stratified vs. random training set allocation on prediction accuracy

Analysis of residual squares comparing stratified and randomly allocated training sets showed that prediction performance varied across lodging severity categories.

In the no lodging category (0%), random allocation produced a lower mean residual square (88.6) than stratified sampling (254.0), although both approaches included a small number of plots with residual squares of approximately 2300-3300 (Fig 12 and Table S8). Similarly, for low lodging severities (5-30%), random allocation consistently produced lower mean residual squares (111.1-216.1) than stratified sampling (231.0-309.1), while a small number of plots under both approaches had residual squares ranging from approximately 1600-4100 (Fig 12, Table S8).

**Fig 12.**
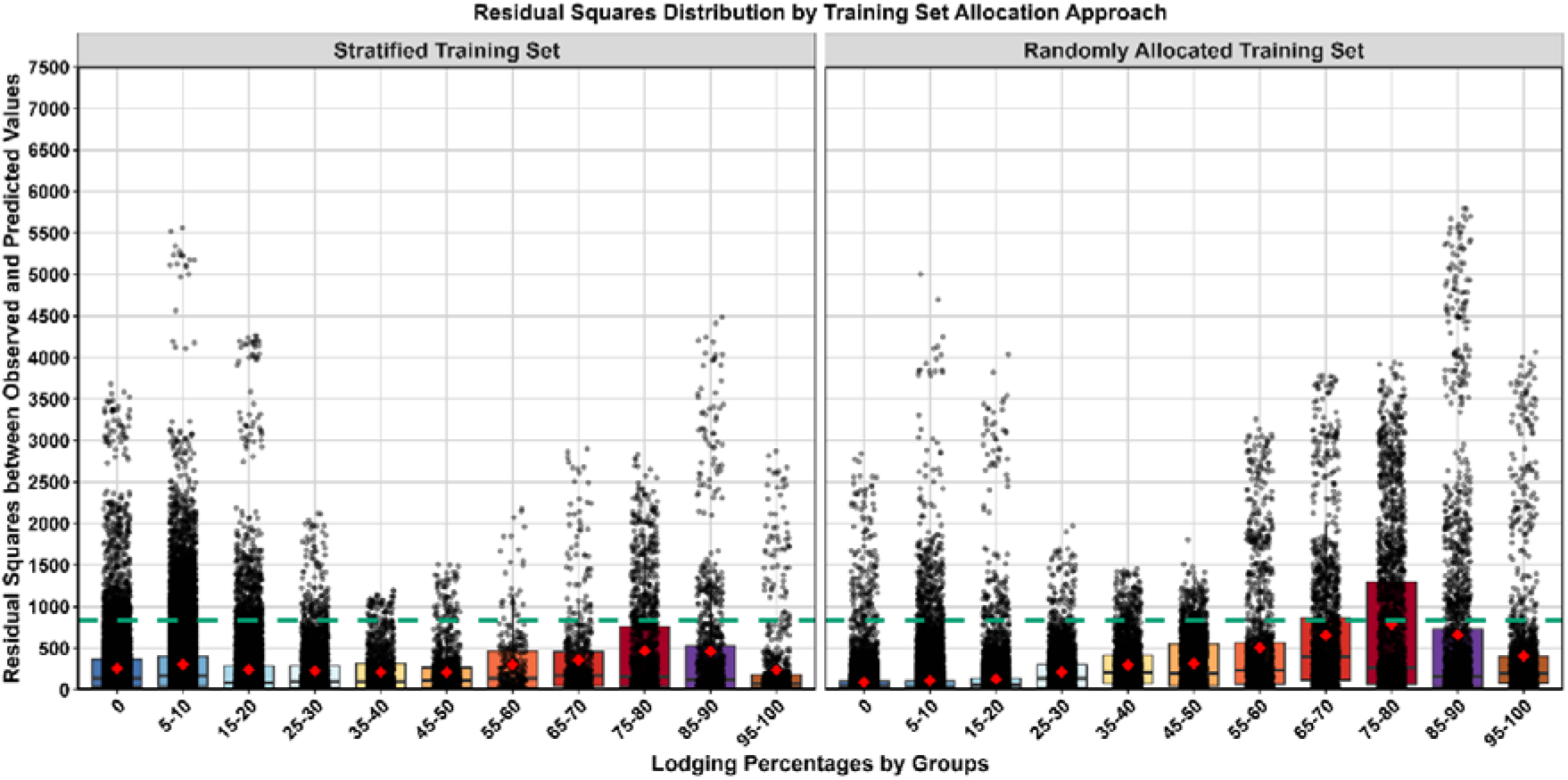
Distribution of residual squares across different lodging severity categories, comparing stratified and randomly allocated training approaches.

In moderate lodging categories (35-60%), stratified sampling resulted in lower mean residual squares than random allocation. Mean residual squares were 223.6 versus 294.2 for 35-40% lodging group, 209.3 versus 326.4 for 45-50% lodging group, and 305.9 versus 490.4 for 55-60% lodging group for stratified and random allocation (Table S8).

A similar pattern was observed for high lodging categories (65-80%). Mean residual squares under random allocation were 642.2 and 806.3 for the 65-70% and 75-80% lodging categories compared with 346.6 and 485.1 under stratified sampling (Table S8). These values corresponded to approximately 85% and 66% higher mean residual squares under random allocation. Random allocation also showed broader residual distributions in these categories, with some plots reaching residual squares of approximately 2,700-3,600 (Fig. 12).

For severe lodging categories (85-100%), stratified sampling continued to produce lower mean residual squares. In the 85-90% category, the mean residual square was 436.8 under stratified sampling compared with 675.4 under random allocation. Similarly, for the 95– 100% lodging category, the mean residual square was 234.4 under stratified sampling compared with 400.3 under random allocation (Table S8).

Overall, random allocation performed better only in low lodging categories (0-30%) with 15-72% lower mean residual squares, while stratified sampling demonstrated superior performance across moderate to severe lodging categories (35-100%), with advantages ranging from 24-85%.

## 4. DISCUSSION

The aim of this study was to evaluate whether percentile height data derived from UAV-enabled photogrammetry could be used to quantify lodging severity in sorghum breeding trials. Our approach focused on detecting lodging by examining relative changes in percentile height within plots before lodging (flight 1) and after lodging occurred (flight 2), using geo-referenced height measurements extracted from UAV point clouds. A range of predictive modelling techniques was implemented, including parametric and non-parametric algorithms as well as a naïve ensemble method. Model performance was evaluated across different lodging severity levels and sites. We also investigated the influence of training-set composition by comparing random and stratified allocation approaches across the lodging severity distribution. In addition, poorly predicted plots were examined through visual inspection of UAV-derived plot images to identify potential sources of disagreement between manual and UAV-predicted lodging severity.

Across these analyses, our ensemble model demonstrated moderate to strong Pearson correlations (0.73 to 0.84) between manual lodging counts and UAV-predicted counts. The comparison of training-set allocation approaches further showed that random allocation performed better primarily at low lodging severities (0-30%), whereas stratified allocation improved prediction performance across moderate to severe lodging categories (35-100%). Together, these results demonstrate that UAV-derived percentile height changes can be used to quantify lodging severity across a broad range of lodging conditions. This UAV-based approach provides a practical option for sorghum breeding programs by reducing phenotyping costs and increasing the number of experimental plots that can be assessed for lodging severity, thereby increasing the amount of phenotypic information available for selection.

### 4.1 Lodging can be predicted using UAV-derived photogrammetric height data

This study demonstrates that lodging derived from manual counts in sorghum breeding trials can be accurately estimated with reasonable accuracy using UAV-derived photogrammetric height data. The integration of multispectral imagery enables detailed analysis of plot height variations before and after lodging events. While Chauhan et al. (2019) noted that remote sensing methods are still experimental, recent studies have validated UAV-derived sensing metrics for lodging measurements across different crops: maize (Shu et al. 2023), wheat (Wilke et al. 2019; Zhang et al. 2020), and rice (Yang et al. 2017). The findings from this current study confirm the capability of UAV-based imagery to assess lodging severity in sorghum breeding programs across sites through photogrammetric height measurements. This research, along with previous studies, supports UAV imagery as an essential tool for assessing lodging in breeding programs, particularly important given lodging’s significant impact on crop yield and quality.

### 4.2 Comparative performance of statistical learning approaches for lodging prediction

This study compared parametric, non-parametric, and ensemble models for predicting plot-level lodging severity from UAV-derived photogrammetric height data. Given that predictors quantified plot-level changes in canopy height, substantial nonlinear responses were not expected. The results for the evaluation metrics largely supported this, with all models producing reasonable results with RMSE values of 17 to 18 percentage points, r values above 0.80, and R² values generally greater than 60%. The RMSE values are affected by the plots that have high residual squares.

The parametric models, which depend solely on linear assumptions, consistently produced lower predictive accuracy than the non-parametric models. This suggests that percentile-based height metrics captured additional structural variation in the canopy that was not fully represented by mean height alone, enabling non-parametric models to exploit these relationships and improve prediction accuracy (Ryo & Rillig, 2017). Such metrics likely reflect variation in traits such as absolute plant height and senescence, which are physiologically linked to lodging susceptibility (Wang et al. 2022). A similar superior performance of non-parametric models has been observed in recent studies utilizing a UAV-derived sensing matrix to predict lodging susceptibility in crops such as wheat and soybean (Sarkar et al. 2023; Wang et al. 2022; Yang et al. 2017).

Our research showed differences in performance between parametric and non-parametric models, due to the distinctive features underlying their predictions. Despite their different approaches, no single parametric approach was consistently superior to the others, and similarly, no non-parametric model was consistently superior to the other models of this type. This indicated that each model was making use of different features in the data, and their performance varied based on the specific datasets used, and no single model consistently produced the best predictions. Similarly, multiple studies have highlighted the challenge of identifying a single “best” individual model, illustrating that some models perform well in particular contexts but are not universally superior (Merrick & Carter 2021; Messina et al. 2025a; Plavšin et al. 2022; Tomura et al. 2025). Therefore, an ensemble approach was employed by averaging results from all of the models, resulting in a naïve average that weighted each prediction algorithm equally.

This ensemble naïve average model was performed slightly better than both the parametric and non-parametric individual models across all evaluation metrics. These findings align with recent research on UAV-based phenotyping (Shu et al. 2022; Zang et al. 2024), physiological trait modelling (Tanaka et al. 2024), and lodging detection (Ali et al. 2023). The superior performance of ensemble naïve averaging can be attributed to its ability to leverage distinct model strengths while reducing individual weaknesses (Bruno et al. 2022; Rokach 2010). The improved performance of ensemble naïve averaging can be theoretically explained through the diversity prediction theorem (Page, 2018), which demonstrates that ensemble effectiveness depends on the ability of constituent models to extract different features from the data rather than simply averaging prediction errors. Our findings support this framework, as prediction diversity observed among individual models contributed to reduced ensemble error, resulting in higher prediction accuracy. By combining predictions from multiple models, this approach effectively reduces individual model biases, resulting in more robust predictions. This advantage is particularly evident in scenarios where single models may have limited predictive capabilities. This aligns with recent genomic prediction studies showing that information diversity among individual models critically influences ensemble performance (Tomura et al. 2025).

Empirical studies consistently demonstrate that ensemble methods achieve higher accuracy and lower error rates compared to individual models (Bonab & Can 2019; Bruno et al. 2022; Messina et al. 2025; Tomura et al. 2025). The ensemble approach demonstrates the effectiveness of multi-model integration for lodging estimation in phenotyping applications, offering a robust framework that addresses phenotypic variability and supports breeding program advancement.

### 4.3 Investigation of naïve ensemble model performance across lodging severity levels

The naïve ensemble model showed differences in prediction accuracy and prediction error across lodging severity levels. Moderate to high prediction accuracies were observed for low to moderate lodging severities (0-60%) and at severe lodging (95-100%) (Fig 8). Previous studies have similarly reported that UAV-based methods can distinguish phenotypic differences at the lower and upper ends of lodging severity range, whereas prediction accuracy prediction can be more difficult at intermediate lodging severities because visual and canopy characteristics may overlap among lodging categories (Singh et al. 2019; Sun et al. 2019). UAV imagery methods provided more accurate estimates of lodging severity and better captured variability within the moderate range (15-60%) (Zang et al. 2024).

To further understand the sources of prediction error, 180 poorly predicted plots were identified from the complete dataset of 2,675 plots based on residual squares (Fig. 8; Table S4). These plots represented 6.8% of the complete dataset. Plot-level images extracted from the UAV imagery were visually compared with the manually measured and UAV-predicted lodging counts. This assessment identified two main factors associated with poor prediction performance: potential transcription or recording errors in the manual lodging data and poor plot establishment.

The first factor was potential transcription or recording errors in the manual lodging data. For example, some plots showed little or no visible lodging in the single-plot UAV images despite having high recorded manual lodging counts, including approximately 90% lodging (Figs. S9 and S10). In contrast, other plots with approximately 90% manually measured and UAV-predicted lodging showed stalks clearly lying on the ground in the UAV images (Figs. S11 and S12). These contrasting examples indicated that some of the large differences between the manually measured and UAV-predicted lodging values were likely associated with the recorded manual phenotypic data rather than errors in the plant-counting procedure itself.

Overall, 82 of the 180 poorly predicted plots showed potential transcription or recording errors in the manual lodging data (Table S4). These plots represented 45.6% of the poorly predicted subset but only 3.1% of the complete dataset. Visual inspection indicated that the recorded manual lodging counts for these plots did not correspond with the lodging condition visible in the UAV-derived single-plot images (Figs. S9-S13). These discrepancies were therefore considered likely to have resulted from transcription, recording, or plot-assignment errors during field phenotyping and data entry. Previous studies have similarly reported discrepancies associated with manual lodging assessment (Sarkar et al. 2023; Singh et al. 2019).

Prediction accuracy decreased at higher lodging severities of 65-80%. This reduction may have been partly associated with potential errors in the recorded manual lodging data and the greater difficulty of distinguishing differences among intermediate to highly lodged plots based on changes in canopy structure (Chauhan et al. 2019; Liu et al. 2023). However, prediction accuracy increased again at severe lodging levels of 95-100% compared with the 85-90% range. At these severe lodging levels, plants lying close to the ground produce larger changes in canopy height, which may provide a clearer signal for prediction using UAV-derived photogrammetric height data (Yang et al. 2017).

The second factor associated with poor prediction was poor plot establishment. Poor establishment was observed in 29 of the 180 poorly predicted plots, representing 16.1% of the poorly predicted subset and 1.1% of the complete dataset (Table S4). These plots had establishment ratings of 4-6 and occurred more frequently among the poorly predicted plots than expected based on their proportion in the complete dataset. Poorly established plots have fewer plants and greater exposure of the ground surface, which can influence plot-level percentile height measurements and consequently affect height-based lodging predictions. Similar effects have been reported in UAV phenotyping, where poor crop establishment and increased exposure of bare ground can interfere with the estimation of canopy characteristics (Morris et al. 2022; B. Zhao et al. 2018). More importantly, measurements of grain yield, plant height, flowering, lodging, and other traits from poorly established plots may not accurately represent the genetic performance of the tested material. Therefore, such plots provide limited value for selection in breeding trials.

For the remaining poorly predicted plots, no obvious cause of prediction error was identified for 59 plots, five were associated with errors in the UAV-based lodging estimates, and plot-level images could not be extracted for visual inspection for another five plots (Table S4). The absence of an obvious cause for some poorly predicted plots may reflect complex canopy conditions, including plants leaning against one another or lodging patterns that were difficult to distinguish from single-plot UAV images. Similar challenges associated with complex canopy structures have been reported in UAV-based phenotyping (Liu et al. 2017).

### 4.4 Consistency of performance of ensemble model

Our research systematically removed manual measurement errors and poor-establishment plots to develop a robust plot-level lodging prediction model for sorghum breeding trials. Models retrained on these refined datasets revealed that ensemble, RF, and SVM radial approaches exhibited similar performance metrics (Fig S10). This convergence is expected under the ensemble error-diversity principle: ensembles provide the greatest benefit when individual models are differentially affected by problematic observations. In the full dataset, some methods were more sensitive to high-error plots, and the ensemble improved average performance by compensating for these method-specific failures. After excluding extreme-error plots, the remaining data were more coherent, reducing the incremental advantage of the ensemble-based model as expected. This result suggests that removing known high-error plots reduced the complex patterns in the data, allowing RF and SVM radial to perform consistently with the ensemble model. This result is consistent with previous findings (Gigović et al. 2019; Sharda et al. 2025), which found that noise reduction through excluding known errors improved model performance and reduced the gap between complex and simpler algorithms. Despite this, we selected the ensemble model for further analysis as it demonstrated greater resilience to errors and noisy data when known error plots were included in the training data (Fig 5), confirming its advantage in handling problematic data. The robustness of our ensemble model is consistent with prior research demonstrating that ensemble approaches leverage model diversity to mitigate noise impacts and enhance predictive reliability in challenging data scenarios (Ferhi et al. 2025; Xu et al. 2020).

Quantitative comparison of five analytical approaches (Fig 7) revealed differential contributions of each error source to overall model performance. Removing plots with manual measurement errors reduced prediction error by 13.8% compared with the original model, while excluding plots with poor establishment alone reduced error by 3.5%. The substantially larger impact of manual measurement errors likely reflects that these represent fundamental misclassifications in ground truth data, which directly confound model learning. In contrast, poor establishment introduces error primarily through altered image features (increased bare ground visibility) rather than incorrect labels. Excluding both error-causing plot types resulted in an 18.5% reduction in prediction error. The retrained model, developed after removing both error sources, demonstrated a 19% lower error than the original model. However, comparison between the approach with both errors removed and the retrained model showed less than 1% improvement from retraining, suggesting that error exclusion alone captures most potential performance gains, and re-training of the model is not necessarily needed.

Manual measurement errors represent the primary source of prediction error in lodging assessment, contributing approximately four times more error than poor establishment. This finding highlights that investment in accurate manual measurement provides substantially greater returns for breeding programs. While poor establishment contributes modestly to overall error (3.5%), its exclusion remains justified because these plots fundamentally compromise lodging estimation accuracy and are typically excluded from breeding selection decisions. The minimal additional improvement from retraining (<1%) indicates that careful data curation through systematic error identification and exclusion may be more efficient than iterative model retraining for achieving robust lodging predictions in breeding programs.

The differing patterns in prediction accuracy between sites (Fig 8) are driven by plots that were poorly predicted despite having no identifiable cause. Additionally, the number of plots in each lodging severity category within a site also affects prediction accuracy. Except for these factors, our study did not identify any differences between sites.

Training set design had minimal influence on model performance: stratified sampling improved predictions for high lodging counts (65-90%), while random allocation favoured low lodging prediction (0-30%), reflecting the dataset’s natural predominance of low-lodging plots (54.2% of plots had between 0-15% lodging). Model performance for moderate lodging predictions (30-60%) remained stable across both approaches. The causes for these differences are complex and difficult to specifically identify but are likely caused by the following reasons.

### 4.5 Application of UAV height data derived from photogrammetric approaches for high-throughput assessment of observed lodging in sorghum breeding trials

Our research showed that UAV-derived multispectral photogrammetry effectively predicted lodging in sorghum breeding trials, offering advantages in implementation, scalability, cost-effectiveness, and environmental adaptability. This approach aligns with the findings of Maimaitijiang et al. (2020), demonstrating accurate canopy height measurements using RGB photogrammetric point clouds. Wilke et al. (2019) further validated this method’s reliability by showing high correlation coefficients in quantifying lodging percentage and severity in wheat trials. Alternative approaches using artificial intelligence (AI) infused image analysis provide potential approaches to achieve greater accuracy but require extensive training datasets across diverse environments and germplasm (Jiang & Li 2020; Yang et al. 2020). The proposed framework developed here is not only operationally much simpler than other more computationally intensive AI-driven approaches but also demonstrates agility in balancing accuracy with practicality, enabling efficient lodging estimation and monitoring in sorghum breeding plots. Therefore, a UAV-enabled photogrammetry approach is likely to provide breeding programs with a robust, practical solution for making data-driven selections on lodging in sorghum based on accurate, timely data.

## 5. CONCLUSION

This study demonstrates that lodging can be accurately predicted using photogrammetry by analysing changes in percentile heights before and after lodging. The ensemble naïve-average model achieved slightly higher r and R² values and lower RMSE than the individual models, indicating superior overall performance. Although plots with incorrect manual measurements and poor establishment reduced prediction accuracy, the ensemble model remained robust, maintaining strong predictive performance even when such plots were present in the training data.

An important direction for future research is to validate the developed framework at an independent site that was not used for model development. Future work should also evaluate the utility of UAV-predicted lodging counts at the genotype level by accounting for the field experimental design and comparing UAV-predicted and manually measured lodging counts using heritability estimates, genotype rankings and selection decisions. These analyses would provide a more comprehensive assessment of the value of UAV-predicted lodging for identifying lodging-resistant sorghum genotypes in breeding trials.

## ACKNOWLEDGMENTS

We would like to acknowledge The University of Queensland (UQ), and the Queensland Alliance for Agriculture and Food Innovation (QAAFI), as well as the Queensland Government Department of Primary Industry (DPI) and the Grains Research and Development Corporation (GRDC) for providing access to the data used in this research. I would like to thank Shunichiro Tomura for his help his. His research and presentations on the ensemble approach really helped me understand the concept better and learn more about it.

## DATA AVAILABILITY

The data supporting the findings of this study are available from the corresponding author upon reasonable request.

## CODE AVAILABILITY

The R code used for data analysis and model implementation in this study is publicly available at https://github.com/SrinivasaMothukuri/Statistical-analysis-for-lodging-prediction.

## SUPPLEMENTARY MATERIALS

Supplemental material includes Figs. S1 to S19 and Tables S1 to S8. These materials provide additional information on the UAV image-processing workflow, data cleaning and quality assessment, representative plot-level percentile heights, cross-validation design, lodging distributions, predictor selection, model-performance evaluation, identification and assessment of plots with large prediction errors, cross-environment prediction, site-specific residual patterns, comparison of random and stratified sampling approaches, and spatial comparisons of manually observed lodging, UAV-predicted lodging, and plot-level prediction errors.

## ABBREVIATIONS

AI: artificial intelligence
AYTF: advanced yield trial for females
AYTM: advanced yield trial for males
DPHI: difference in percentile height index
DPI: Department of Primary Industries
GCP: ground control point
GNSS: global navigation satellite system
GRDC: Grains Research and Development Corporation
IQR: interquartile range
KNN: k-nearest neighbours
LAI: leaf area index
LGD: lodging
MLR: multiple linear regression
NAS: network-attached storage
OLS: ordinary least squares
PCA: principal component analysis
PCC: Pearson correlation coefficient
PHI: percentile height index
PLS: partial least squares
PLSR: partial least squares regression
QAAFI: Queensland Alliance for Agriculture and Food Innovation
RBF: radial basis function
RF: random forest
RMSE: root mean square error
RR: ridge regression
SVM: support vector machine
UAV: unmanned aerial vehicle
UQ: University of Queensland
USA: United States of America
Vcmax: maximum carboxylation capacity.

