## Supplementary material for "Phenomic Prediction I: Plot-Level Prediction of Lodging Severity in Sorghum Breeding Trials Using UAV-Based Photogrammetric Height Data": Not available: Supplementary_Information.pdf (from another version of this preprint: version 1, 2026-03-30)

### Supplementary Information (SI)

### A Framework for Predicting Lodging Severity in Sorghum Breeding Trials Using UAV-Based Photogrammetrically Derived Height Data

Srinivasa R. Mothukuri[1], Sean R. Massey-Reed[1], Andries B. Potgieter[3], Kenneth Laws[2], Colleen Hunt[1,2], Esinam N. Amuzu-Aweh[1], Mark Cooper[3,5], Emma Mace[1,2], David Jordan[1]

**Affiliation:**

[1] The University of Queensland, Queensland Alliance for Agriculture and Food Innovation (QAAFI), Hermitage Research Facility, Warwick, QLD, Australia

[2] Agri-Science Queensland, Department of Primary Industries (DPI), Hermitage Research Facility, Warwick, QLD, Australia

[3] The University of Queensland, Queensland Alliance for Agriculture and Food Innovation (QAAFI), St Lucia, QLD, Australia

[4] The University of Queensland, Queensland Alliance for Agriculture and Food Innovation (QAAFI), Gatton, QLD, Australia

[5]Australian Research Council Centre of Excellence for Plant Success in Nature and Agriculture, The University of Queensland, Brisbane, QLD, Australia

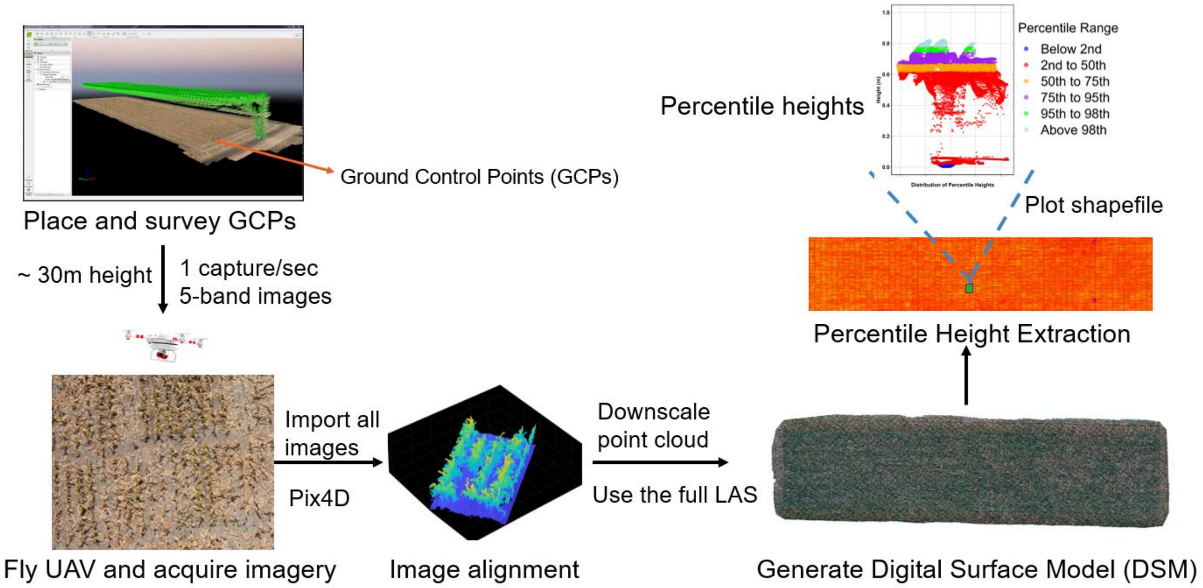


**Fig. S1:** Simplified workflow of UAV imagery pre-processing to extract the percentile heights per plot.

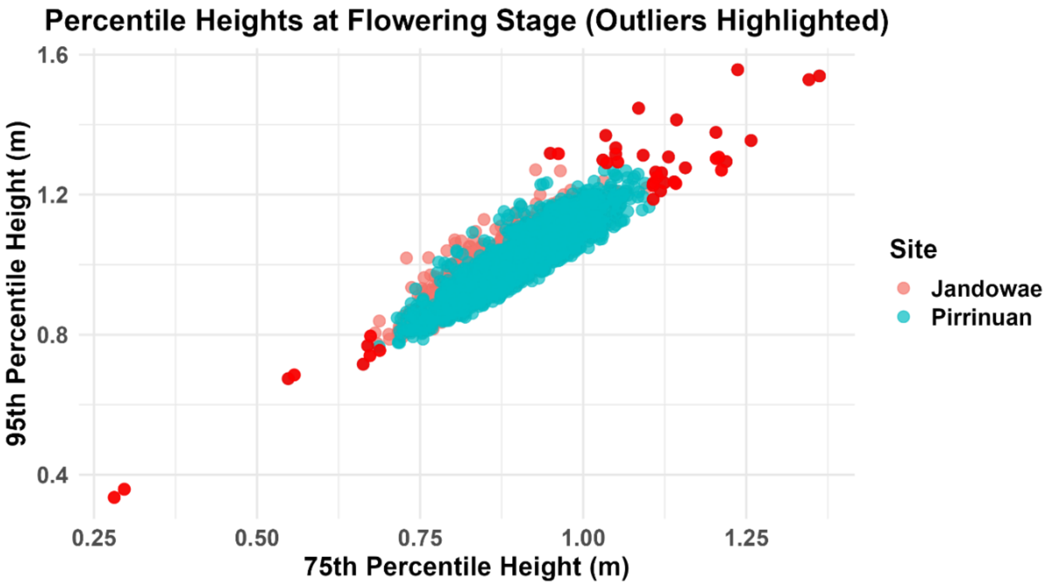


**Fig. S2:** Percentile height at flight -1 (flowering stage) between 75th and 95th percentile by site, highlighting outliers with red dots.

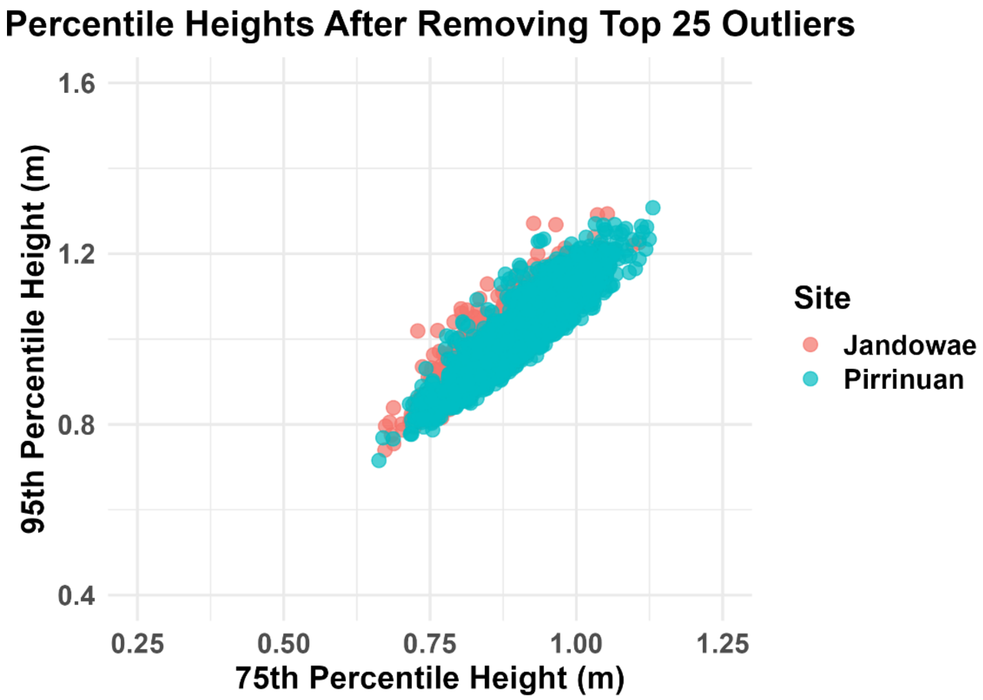


**Fig. S3:** Percentile height after removing top 25 outliers at flight -1 (flowering Stage) between 75th and 95th percentile by site.

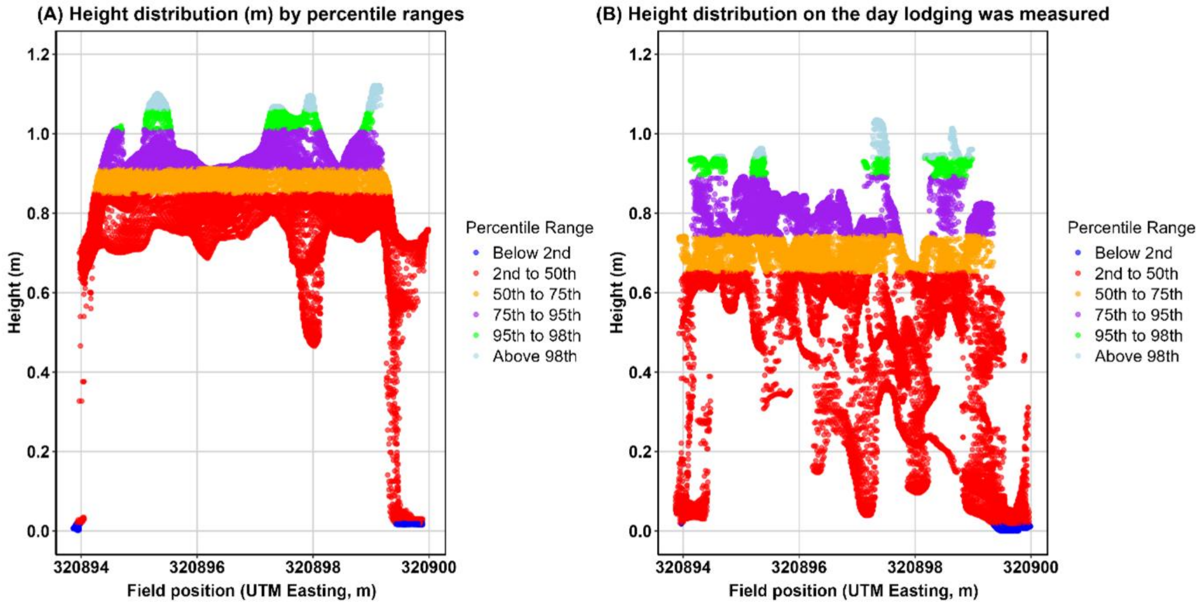


**Fig. S4:** Height distributions for a representative plot with manual lodging severity = 0%. (A) Height measurements around the flowering stage plotted against field position (UTM Easting, m), with points color-coded by percentile ranges. (B) Height measurements for the same plot plotted against the same field position on the date lodging was assessed.

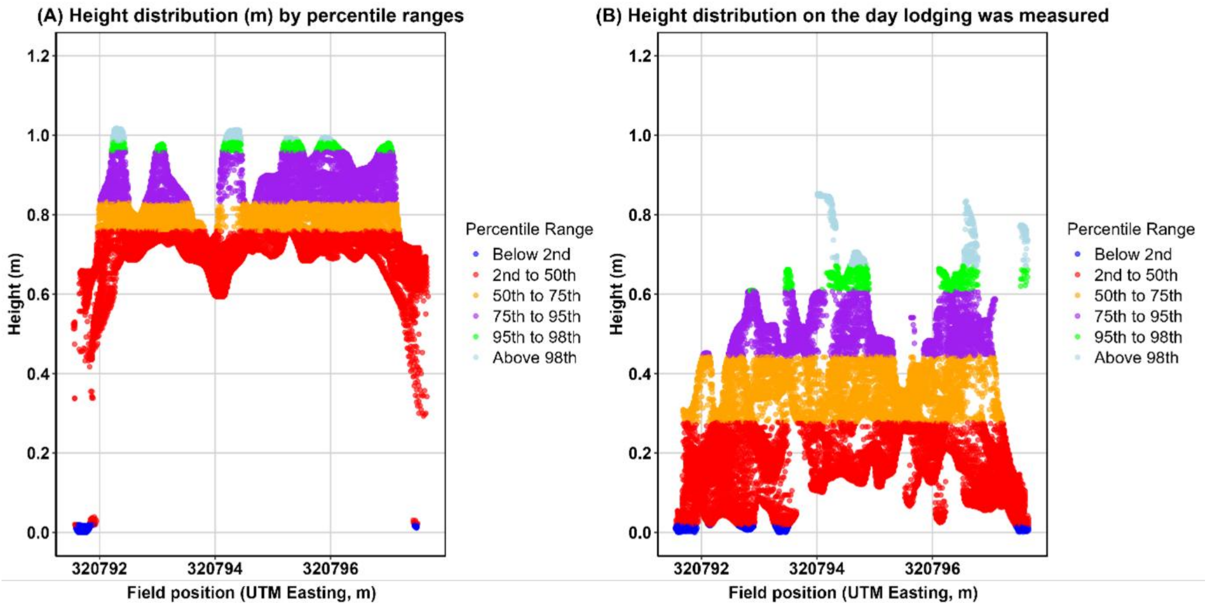


**Fig. S5:** Height distributions for a representative plot with manual lodging severity = 60%. (A) Height measurements around the flowering stage plotted against field position (UTM Easting, m), with points color-coded by percentile ranges. (B) Height measurements for the same plot plotted against the same field position on the date lodging was assessed.

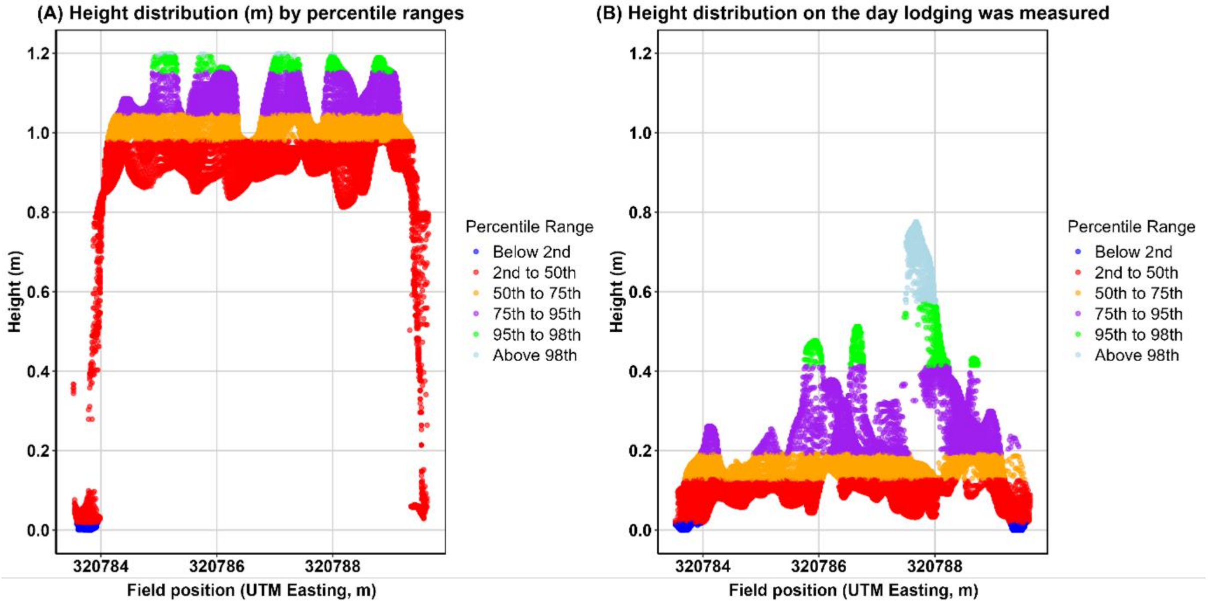


**Fig. S6:** Height distributions for a representative plot with manual lodging severity = 90%. (A) Height measurements around the flowering stage plotted against field position (UTM Easting, m), with points color-coded by percentile ranges. (B) Height measurements for the same plot plotted against the same field position on the date lodging was assessed.

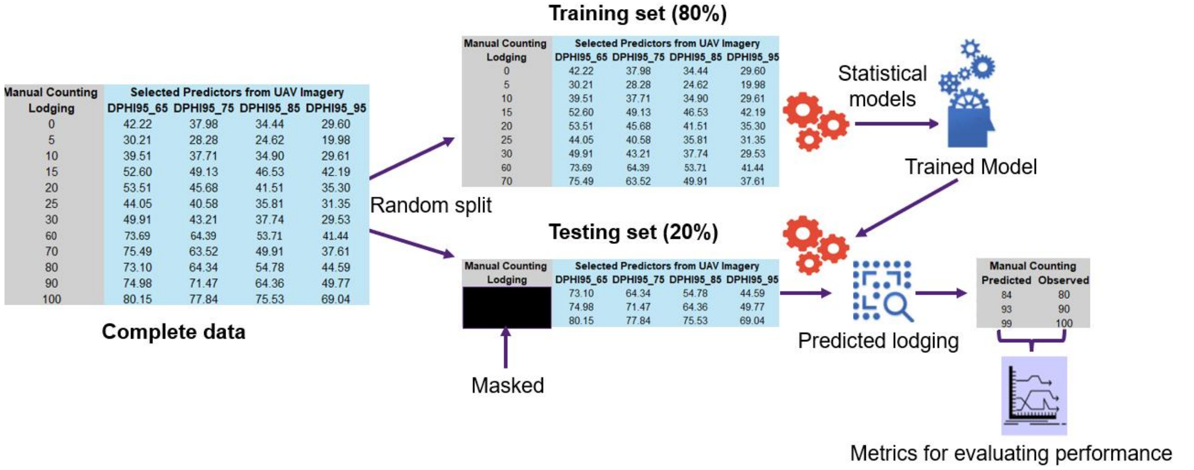


**Fig. S7:** Schematic representation of the cross-validation workflow for lodging prediction using UAV imagery data with 80-20 train-test split validation.

| Percentages | 0 | 5-10 | 15-20 | 25-30 | 35-40 | 45-50 | 55-60 | 65-70 | 75-80 | 85-90 | 95-100 | Total |
|---|---|---|---|---|---|---|---|---|---|---|---|---|
| No. Plots | 625 | 829 | 336 | 160 | 103 | 105 | 69 | 101 | 146 | 114 | 87 | 2675 |

**Table. S1:** Distribution of plots by lodging percentages across specified ranges

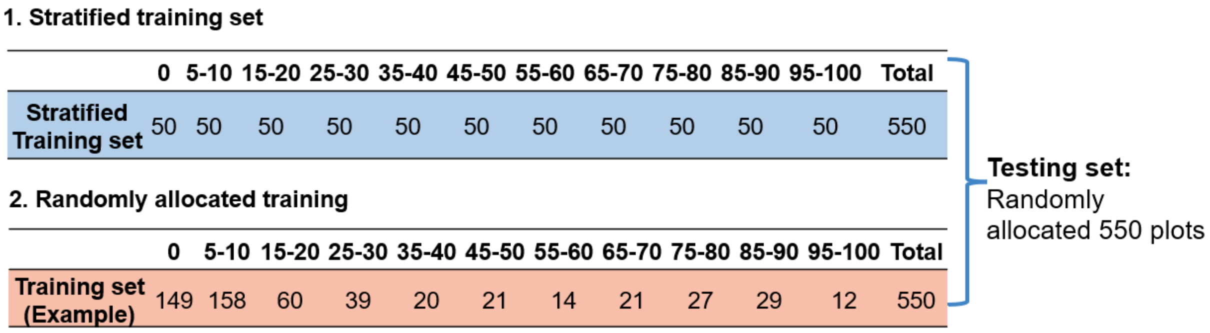


**Fig. S8:** Comparison of stratified and random training data allocation schemes for lodging severity classification with 550 plots and 100 iterations. The random training set is an example of the random allocation of plots into training and testing sets.

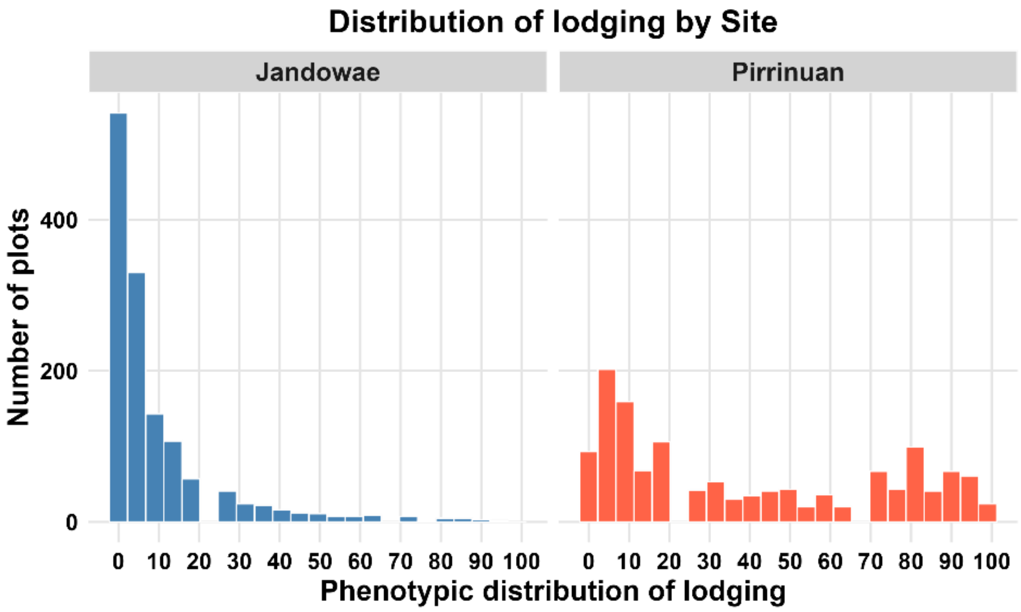


**Fig. S9:** Distribution of phenotypic lodging scores by site (Jandowae and Pirrinuan).

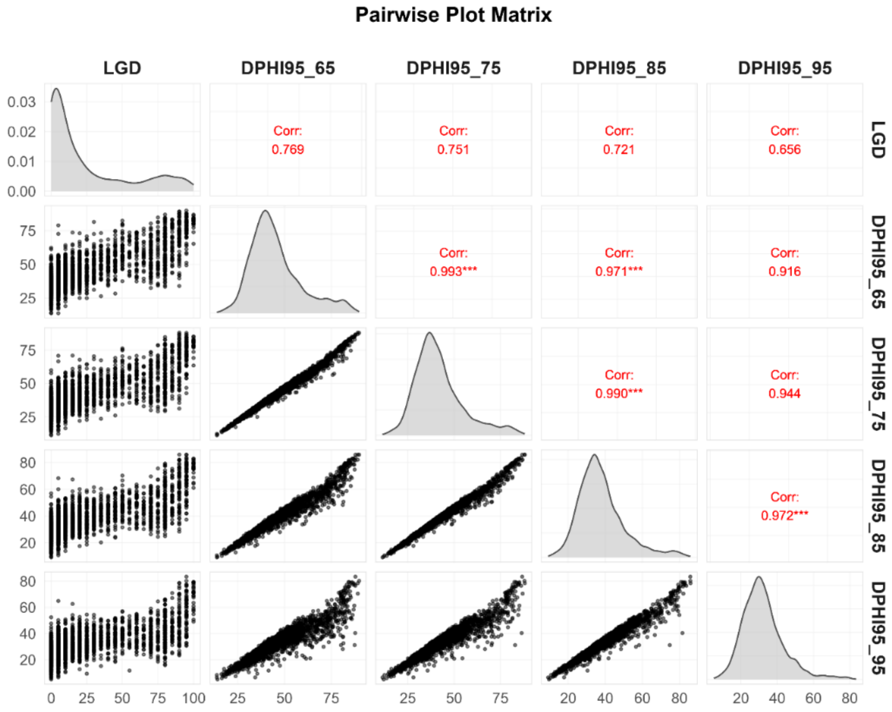


**Fig. S9:** A pairwise plot matrix showing relationships between several variables: manual lodging (LGD) and DPHI95 at different levels (65, 75, 85, and 95).

**Table. S2:** Comparison of performance metrics between parametric and non-parametric models across 100 iterations for lodging prediction using cross-validation approach; correlation coefficients (r, $R^2$) and residual mean square error (RMSE) statistics across 100 model iterations.

| Model | Min r | Max r | Mean r | Min r2 | Max r2 | Mean r2 | Min RMSE | Max RMSE | Mean RMSE |
|---|---|---|---|---|---|---|---|---|---|
| **MLR** | 0.74 | 0.82 | 0.78 | 0.54 | 0.68 | 0.61 | 16.75 | 19.99 | 18.54 |
| **RR** | 0.74 | 0.82 | 0.78 | 0.54 | 0.68 | 0.61 | 16.82 | 19.96 | 18.55 |
| **PLSR** | 0.74 | 0.82 | 0.78 | 0.54 | 0.68 | 0.61 | 16.81 | 20.05 | 18.58 |
| **KNN** | 0.74 | 0.83 | 0.79 | 0.54 | 0.69 | 0.63 | 16.08 | 19.98 | 18.16 |
| **RF** | 0.74 | 0.84 | 0.80 | 0.55 | 0.70 | 0.63 | 16.19 | 19.84 | 17.98 |
| **SVM** | 0.74 | 0.84 | 0.80 | 0.55 | 0.70 | 0.64 | 16.10 | 20.41 | 18.25 |
| **Ensemble** | 0.75 | 0.84 | 0.80 | 0.56 | 0.70 | 0.64 | 16.01 | 19.54 | 17.89 |

**Table S3:** Mean residual, residual squares and ratio below statistical threshold (residual squares above 1282) across lodging severity categories.

| Lodging percentages | Plots | Mean Residual Squares | Ratio below statistical threshold |
|---|---|---|---|
| 0 | 625 | 130 | 98.9% |
| 5-10% | 829 | 145 | 98.7% |
| 15-20% | 336 | 175 | 97.5% |
| 25-30% | 160 | 254 | 97.4% |
| 35-40% | 103 | 286 | 100% |
| 45-50% | 105 | 313 | 92.4% |
| 55-60% | 69 | 569 | 76.7% |
| 65-70% | 101 | 1243 | 60.8% |
| 75-80% | 146 | 1363 | 56.3% |
| 85-90% | 114 | 976 | 78.6% |
| 95-100% | 87 | 325 | 95.8% |

**Table S4:** Visual inspection using single plot UAV images on plots above statistical threshold (residual squares above 1282) to identify the cause of error.

| Error category | Number of plots (n) | Percentage (%) |
|---|---|---|
| Manual scoring error | 82 | 45.56% |
| Poor establishment | 29 | 16.11% |
| No obvious cause identified | 59 | 32.78% |

| | | |
|---|---|---|
| UAV estimate error | 5 | 2.78% |
| Single-plot image for visual inspection was not extracted | 5 | 2.78% |
| Total | 180 | 100% |

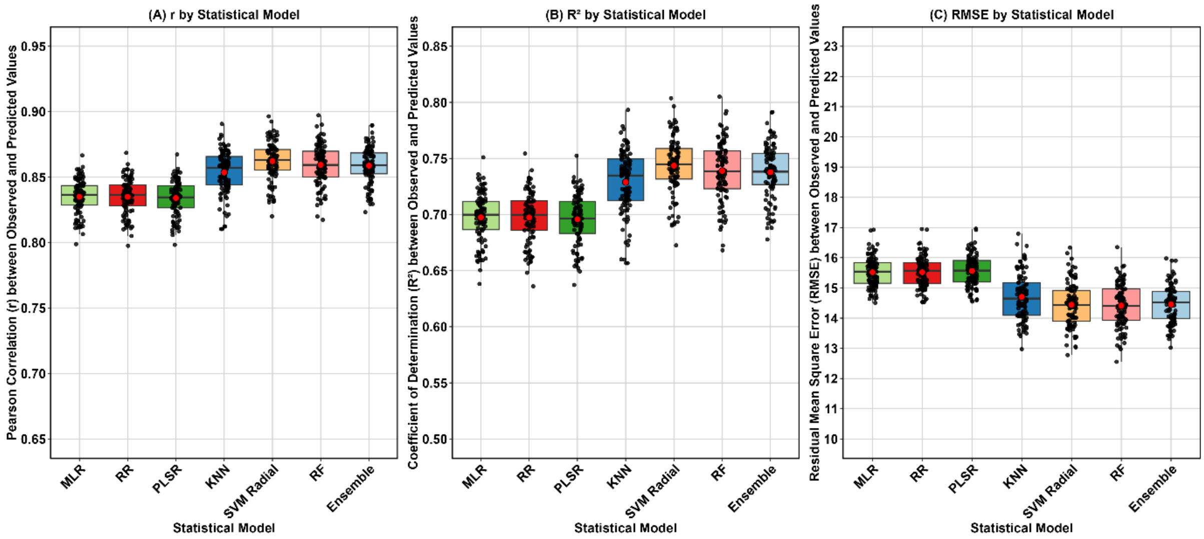


**Fig. S10:** Retrained model evaluation of different statistical models based on r, $R^2$, and RMSE. Black dots illustrate model performance distribution across 100 iterations; red dots indicate mean values.

**Table. S5:** Retrained model performance after excluding known error plots: comparison of parametric and non-parametric models across 100 iterations for lodging prediction.

| Model | Min r | Max r | Mean r | Min $R^2$ | Max $R^2$ | Mean $R^2$ | Min RMSE | Max RMSE | Mean RMSE |
|---|---|---|---|---|---|---|---|---|---|
| **MLR** | 0.80 | 0.87 | 0.84 | 0.64 | 0.75 | 0.70 | 14.53 | 16.92 | 15.51 |
| **RR** | 0.80 | 0.87 | 0.83 | 0.64 | 0.75 | 0.70 | 14.53 | 16.97 | 15.51 |
| **PLSR** | 0.80 | 0.87 | 0.83 | 0.64 | 0.75 | 0.70 | 14.57 | 16.94 | 15.55 |
| **KNN** | 0.81 | 0.89 | 0.85 | 0.66 | 0.79 | 0.73 | 12.94 | 16.71 | 14.67 |
| **RF** | 0.82 | 0.90 | 0.86 | 0.67 | 0.81 | 0.74 | 12.57 | 16.32 | 14.40 |
| **SVM** | 0.82 | 0.90 | 0.86 | 0.67 | 0.80 | 0.74 | 12.61 | 16.09 | 14.27 |
| **Ensemble** | 0.82 | 0.89 | 0.86 | 0.68 | 0.79 | 0.74 | 13.00 | 15.96 | 14.43 |

**Table. S6:** Comparison of ensemble model prediction accuracy (RMSE) across five approaches.

| Approach | Min RMSE | Mean RMSE | Max RMSE | SD RMSE |
|---|---|---|---|---|
| Original Model | 16.01 | 17.90 | 19.54 | 0.73 |
| Scoring Errors Removed | 13.92 | 15.42 | 17.48 | 0.6 |
| Poor Establishment | 14.94 | 17.27 | 19.09 | 0.77 |
| Both removed | 13.16 | 14.59 | 16.02 | 0.59 |
| Retrained model | 13.02 | 14.46 | 15.98 | 0.62 |

**Table S7:** Mean residual, mean residual squares, and ratio below threshold (833) across lodging severity categories by site

| Lodging | Number of Plots | | Mean Residual Squares | | Ratio below mean RMSE² | |
|---|---|---|---|---|---|---|
| Percentages | Jandowae | Pirrinuan | Jandowae | Pirrinuan | Jandowae | Pirrinuan |
| 0 | 526 | 68 | 69.1 | 220 | 99.8% | 92.6% |
| 5-10% | 469 | 321 | 77.5 | 157 | 98.7% | 96.9% |
| 15-20% | 163 | 154 | 72.1 | 193 | 100% | 95.4% |
| 25-30% | 65 | 85 | 183 | 243 | 100% | 96.5% |
| 35-40% | 38 | 58 | 317 | 292 | 89.5% | 94.8% |
| 45-50% | 23 | 68 | 418 | 289 | 87% | 92.6% |
| 55-60% | 14 | 47 | 582 | 446 | 78.6% | 85.1% |
| 65-70% | 16 | 56 | 346 | 686 | 93.8% | 69.6% |
| 75-80% | 6 | 90 | 67.4 | 802 | 100% | 65.6% |
| 85-90% | 8 | 78 | 430 | 641 | 87.5% | 76.9% |
| 95-100% | 2 | 78 | 1153 | 356 | 50% | 92.3% |

**Table S8:** Mean residual, mean residual squares, and ratio below threshold (1282) across lodging severity categories by stratified training set and randomly allocated training set.

| Lodging | Mean Residual Squares | | Ratio below mean RMSE² | |
|---|---|---|---|---|
| Percentages | Stratified | Random | Stratified | Random |
| 0 | 254 | 89 | 95.1% | 98.8% |
| 5-10% | 309 | 111 | 90.9% | 98.1% |
| 15-20% | 246 | 133 | 93.4% | 98.1% |
| 25-30% | 231 | 216 | 95.3% | 97.3% |
| 35-40% | 224 | 294 | 95.8% | 93.8% |

| | | | | |
|---|---|---|---|---|
| 45-50% | 209 | 326 | 93.4% | 91.2% |
| 55-60% | 306 | 490 | 90.2% | 83.6% |
| 65-70% | 347 | 642 | 90.3% | 76.4% |
| 75-80% | 485 | 806 | 78.1% | 65.6% |
| 85-90% | 437 | 675 | 86% | 75.6% |
| 95-100% | 234 | 400 | 92.5% | 91.2% |

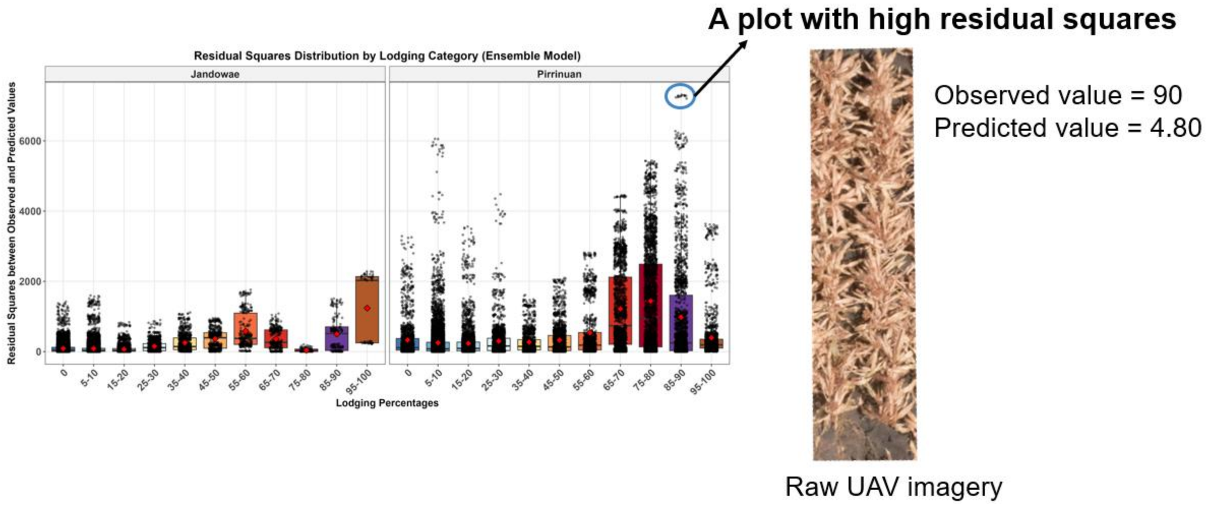


**Fig. S11:** Residual squares distribution by lodging category (ensemble model) across two sites: Jandowae (left panel) and Pirrinuan (right panel). For each site, the x-axis represents different lodging percentage categories (from 0 to 100%), while the y-axis shows the residual squares between observed and predicted values.

The blue circle highlights a case with particularly high error, where there was a large inconsistency between the observed value (90%) and the predicted value (4.80%). The UAV imagery on the right shows this plot contains standing plants with minimal lodging. This indicates the model correctly predicted a low lodging value (4.80%), while the manual observation incorrectly reported a high lodging value (90%), resulting in a very high residual square error. This example highlights a limitation in the manual assessment process rather than an error in the model, as visual evidence from UAV imagery confirms that the model correctly predicted lodging.

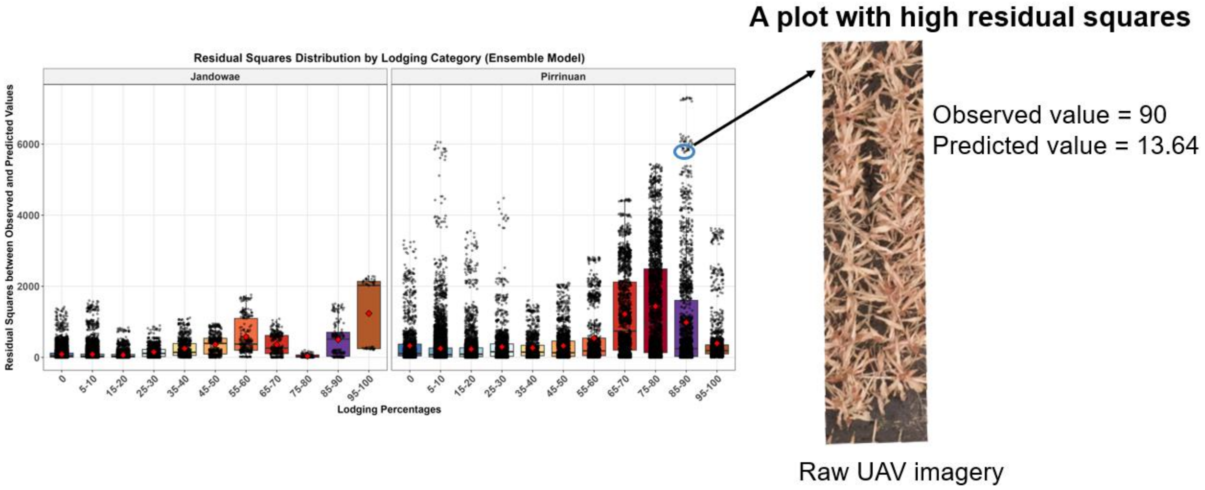


**Fig. S12:** A plot with high residual squares due to an error in manual counting.

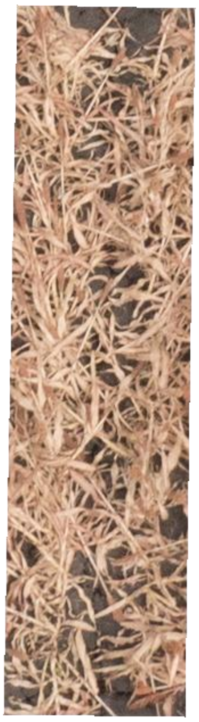

**Fig. S13:** Plot with close to zero residual squares, demonstrating an example between observed and predicted lodging values. The observed value of the plot is 90% lodging, and the model predicted 91% lodging, resulting in a minimal residual error.

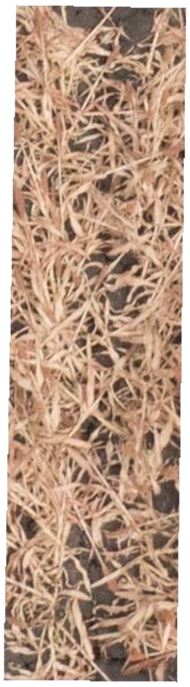

**Fig. S14:** A completely lodged plot, representing an example of matched between observed and predicted lodging values. The observed lodging value is 95% and the model predicted 94.22%, resulting in minimal residual error.

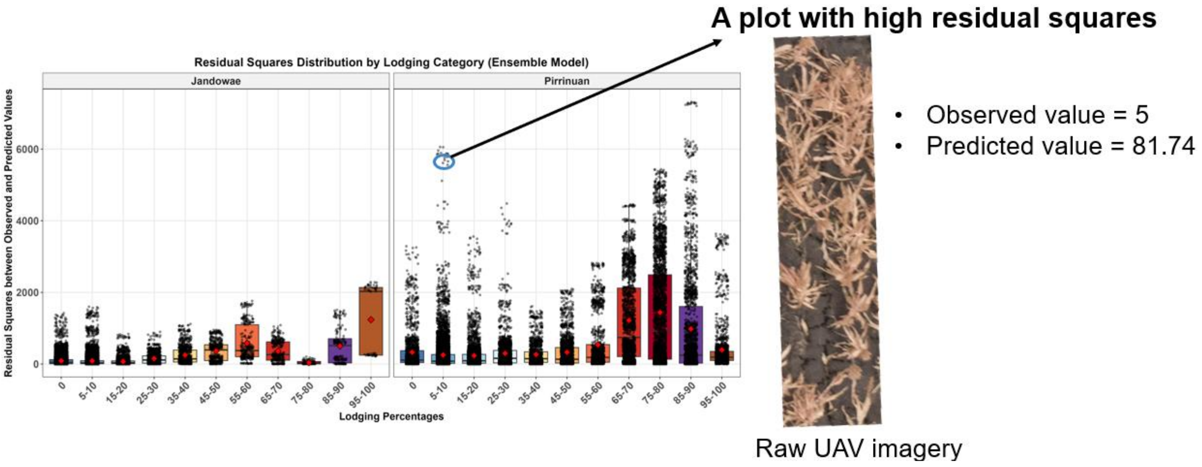


**Fig. S15:** A plot with high residual squares due to poor plot establishment, which can be attributed to water severity and germination rates.

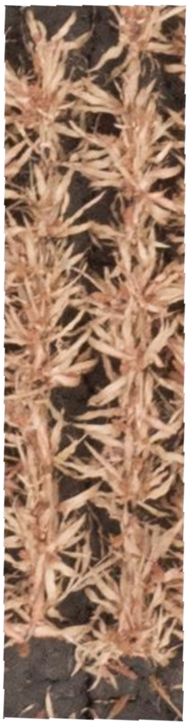

**Fig. S16:** A plot with close to zero residual squares. The observed value of the plot is 0% lodging, and the model predicted 0.1% lodging, resulting in a minimal residual error.
